# Structure-guided synthesis of FK506 and FK520 analogs with increased selectivity exhibit in vivo therapeutic efficacy against Cryptococcus

**DOI:** 10.1101/2022.03.25.485831

**Authors:** Michael J. Hoy, Eunchong Park, Hyunji Lee, Won Young Lim, D. Christopher Cole, Nicholas D. DeBouver, Benjamin G. Bobay, Phillip G. Pierce, David Fox, Maria Ciofani, Praveen R. Juvvadi, William Steinbach, Jiyong Hong, Joseph Heitman

**Affiliations:** Department of Molecular Genetics and Microbiology, Duke University Medical Center, Durham, North Carolina, USA; Department of Immunology, Duke University Medical Center, Durham, North Carolina, USA; Department of Chemistry, Duke University, Durham, North Carolina, USA; Division of Pediatric Infectious Diseases, Department of Pediatrics, Duke University Medical Center, Durham, North Carolina, USA; UCB Biosciences, 7869 NE Day Road West, Bainbridge Island, Washington, USA; Seattle Structural Genomics Center for Infectious Disease (SSGCID), Seattle, Washington, USA; Duke University NMR Center, Duke University Medical Center, Durham, North Carolina, USA

## Abstract

Calcineurin is an essential virulence factor that is conserved across human fungal pathogens including *Cryptococcus neoformans*, *Aspergillus fumigatus*, and *Candida albicans*. Although an excellent target for antifungal drug development, the serine-threonine phosphatase activity of calcineurin is conserved in mammals and inhibition of this activity results in immunosuppression. FK506 (tacrolimus) is a naturally produced macrocyclic compound that inhibits calcineurin by binding to the immunophilin FKBP12. Previously, our fungal calcineurin-FK506-FKBP12 structure-based approaches identified a non-conserved region of FKBP12 that can be exploited for fungal-specific targeting. These studies led to the design of an FK506 analog, APX879, modified at the C22 position that was less immunosuppressive yet maintained antifungal activity. We now report high resolution protein crystal structures of fungal FKBP12 and a human truncated calcineurin-FKBP12 bound to a natural FK506 analog, FK520 (ascomycin). Based on information from these structures and the success of APX879, we synthesized and screened a novel panel of C22-modified compounds derived from both FK506 and FK520. One compound, JH-FK-05 demonstrates broad-spectrum antifungal activity *in vitro* and is non-immunosuppressive *in vivo*. In murine models of pulmonary and disseminated *C. neoformans* infection, JH-FK-05 treatment significantly reduced fungal burden and extended animal survival alone and in combination with fluconazole. Furthermore, molecular dynamic simulations performed with JH-FK-05 binding to fungal and human FKBP12 identified additional residues outside of the C22 and C21 positions that could be modified to generate novel FK506 analogs with improved antifungal activity.

**Significance:** Due to rising rates of antifungal drug resistance and a limited armamentarium of antifungal treatments, there is a paramount need for novel antifungal drugs to treat systemic fungal infections. Calcineurin has been established as an essential and conserved virulence factor in several fungi, making it an attractive antifungal target. However, due to the immunosuppressive action of calcineurin inhibitors, they have not been successfully utilized clinically for antifungal treatment in humans. Recent availability of crystal structures of fungal calcineurin bound inhibitor complexes have enabled the structure-guided design of FK506 analogs and led to a breakthrough in the development of a compound increased for fungal specificity. The development of a calcineurin inhibitor with reduced immunosuppressive activity and therapeutic antifungal activity would add a significant tool to the treatment options for these invasive fungal infections with exceedingly high rates of mortality.

## Introduction

Invasive fungal infections represent a major global health burden. Many of the deadliest fungal infections, such as cryptococcal meningoencephalitis, occur in an increasing immunocompromised patient population receiving organ transplants, suffering from HIV/AIDS, treated with corticosteroids, or undergoing cancer chemotherapy. In fact, *Cryptococcus neoformans* infection is one of the leading causes of death in HIV/AIDS patients and is most common in areas where HIV/AIDS is widespread(1). Mortality rates as high as 70% with *C. neoformans* and another ubiquitous fungal pathogen, *Candida albicans,* present a complicated treatment challenge(2). In many cases, physicians must treat these infections with combination therapies of antifungal drugs due to the lack of fungal-specific drug targets and the associated difficulty in developing antifungal drugs with low toxicity. A rise in drug resistance across pathogenic fungi such as *Candida* sp. and *Aspergillus fumigatus* further exacerbates this issue(3). Within the space of FDA-approved antifungal drugs, clinicians have very limited options. Only 4 main classes of antifungal drugs (polyenes, azoles, pyrimidine analogs, and echinocandins) are approved for treatment of systemic fungal infection and each treatment suffers from issues related to high toxicity, bioavailability, or innate and acquired resistance in many species(4). There is an urgent need to develop new antifungal strategies and identify novel targets to address the increasing global threat of deadly invasive fungal diseases.

Calcineurin signaling in *C. neoformans, A. fumigatus,* and *C. albicans*, as well as in other fungal pathogens, is highly conserved and leads to the activation of virulence genes and proteins that are essential for the organism’s growth at host body temperature, hyphal development, and survival in serum, respectively (5-9). These essential roles in key virulence attributes make calcineurin an excellent target for antifungal drug development (10, 11). Calcineurin is a serine-threonine specific protein phosphatase that is a heterodimer consisting of a catalytic subunit, CnA, and a regulatory subunit, CnB, and is activated in the presence of calcium and calmodulin(12). When activated in many pathogenic fungi, calcineurin dephosphorylates the transcription factor Crz1 and triggers its nuclear translocation and transcription of downstream target genes (13-15). FK506 (tacrolimus) is a naturally occurring calcineurin inhibitor that is a product of several *Streptomyces* species and has potent antifungal activity (16-18). FK506 acts on fungal cells by binding to FK506 binding protein 12 (FKBP12) and forming a complex that binds to and inhibits calcineurin by sterically blocking substrate access to the active site (19, 20). FKBP12 is a member of a larger family of proteins called immunophilins that bind to immunosuppressive molecules and mediate their activity(21).

Calcineurin signaling is conserved in humans and mediates critical pathways involved in growth and proliferation. Additionally, the mechanism of calcineurin inhibition is also conserved via FK506-FKBP12 complex binding to calcineurin(20). In humans, calcineurin is critical for T cell activation and the initiation of immune responses(22). Activated calcineurin dephosphorylates the Crz1 homolog NFAT, which promotes expression of immune response genes, such as the cytokine IL-2 (23, 24). As a result, the potent immunosuppressive activity of FK506 in humans makes it unsuitable to treat patients with fungal infections(25). In fact, due to this robust immunosuppressive activity, FK506 is FDA-approved as a post-transplant drug and is administered to patients to prevent host rejection of a variety of transplanted organs and tissues (26, 27).

FK506 is not the only naturally occurring calcineurin inhibitor and FKBP12-binding molecule. FK520 (ascomycin) is a natural FK506 analog that is modified at the C21 position with an ethyl group replacing the C21 allyl group in FK506(28). This C21 modification of FK506 results in a molecule with similarly potent antifungal activity and modestly reduced immunosuppressive activity(29). In this case, modification of the C21 position introduced a slight, but detectable fungal specificity. Although the antifungal and immunosuppressive activity of FK520 has been well-characterized, few studies have focused on developing fungal-specific variants of FK520(30). The C21 position of FK520 is on the calcineurin-facing region of the molecule and may influence formation of the ternary inhibitory complex. Conformationally, modifications made to the C22 position of FK506 and FK520 could be influenced by the differences at the C21 residue.

Previous studies have sought to develop non-immunosuppressive calcineurin inhibitors for the treatment of fungal infections. Minor modifications to the structure of FK506/FK520 have been shown to significantly alter both its immunosuppressive and antifungal activity. L-685,818 is a non-immunosuppressive analog of FK520 that was shown to maintain antifungal activity via calcineurin inhibition despite having no activity against human calcineurin (17, 31). Structurally quite similar to FK520, L-685,818 differs only in being hydroxylated at the C18 position. L-685,818 binds human FKBP12 identically to FK506 and the C18 and C21 modifications are both exposed on the calcineurin interacting effector surface (32, 33). While the precise molecular mechanism by which L-685,818 disrupts ternary complex formation is still unknown and complex chemical synthesis has limited further study, it has been hypothesized the introduction of the polar hydroxyl group may disrupt a hydrophobic interface that would otherwise dock with calcineurin.

We previously demonstrated that APX879, a C22 acetylhydrazone-modified FK506 analog, exhibits a clear increase in fungal specificity that translates into therapeutic efficacy(34). Structures of fungal FKBP12-FK506 and ternary structures of calcineurin-FK506-FKBP12 from *A. fumigatus*, *C. neoformans*, *Coccidioides immitis*, and *C. albicans* led to the structure-guided design of APX879 (34, 35). Key structural differences in the 80s loop of FKBP12 proximal to the FK506-binding pocket led to fungal-specific differences in the FKBP12-FK506-calcineurin interface. Specifically, fungal Phe88 or mammalian His88 residues of FKBP12 introduced differences in distance to the C22 acetylhydrazone of APX879 and presented a clear opportunity to exploit this for increasing fungal specificity. Moreover, protein-inhibitor complex crystal structures and molecular dynamic (MD) simulations identified multiple residues around the molecule that could enhance fungal specificity(36). Compared to FK506, APX879 exhibits a ∼70-fold reduction in immunosuppression, corresponding to an overall 4-fold increase in fungal specificity determined by its Therapeutic Index score(34). Guided by the success of C22-modification in the previous study, we generated a series of second-generation FK506 analogs as well as the first examples of C22-modified FK520 analogs. Here, we report the first crystal structure of a fungal FKBP12 bound to FK520 that was instrumental in the design of FK520-based analogs. Strikingly, one such FK520 analog described here is non-immunosuppressive *in vivo* and efficacious in multiple models of *C. neoformans* infection. Our current advances in structural work and medicinal chemistry provide further evidence that iterative development of FK506/FK520 analogs can increase fungal specificity and drive these compounds into a therapeutic window that can be utilized to treat invasive fungal infections.

## Results

### *A. fumigatus* FKBP12 bound FK520 X-ray crystal structure reveals conserved binding pocket structure

We previously solved X-ray crystal structures of FKBP12-FK506-calcineurin ternary complexes from divergent human pathogenic fungi(34). FK506 and FK520 are structurally similar but differ at the C21 position where FK520 lacks the terminal alkene of FK506 (**Fig. 1A**). To gain better insight into the interactions between FK520 and fungal FKBP12, we first determined the X-ray crystal structure of *A. fumigatus* FKBP12 bound to FK520 at 1.7Å (PDB: 7U0S). *A. fumigatus* was selected for FKBP12 structure analysis due to the readiness with which protein crystals form and the presence of key conserved fungal residues in the 80s loop. The FKBP12-binding motif of FK506 and FK520 is conserved and thus FK520 binds to *A. fumigatus* FKBP12 identically **(Fig 1B and 1C)**. We have previously shown that the 80s loop of FKBP12 in fungi is a critical region for facilitating the formation of FKBP12-inhibitor-calcineurin complex(34). Phe88 is a conserved residue in many pathogenic fungi that differs from the mammalian His88 residue. Both the C21 and C22 of FK520/FK506 converge on this differential residue within 5 to 8 Å and represent excellent targets for synthetic modification.

**Figure 1.**
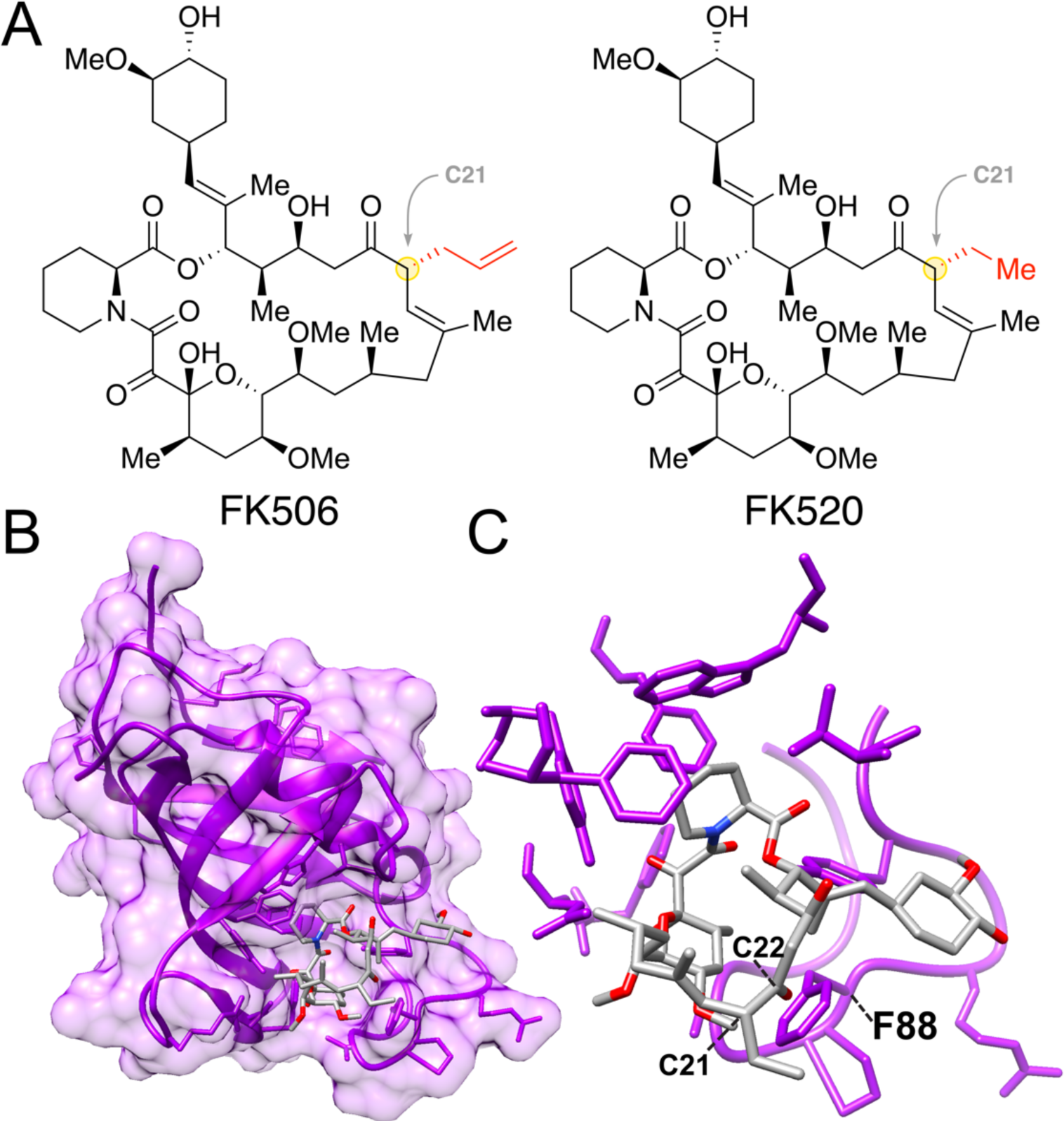
Crystal structure of *Aspergillus fumigatus* FKBP12 bound to FK520. (A) Chemical structures of FK506 (tacrolimus) and FK520 (ascomycin). Structural differences at the C21 residue are highlighted in red. (B) Protein X-ray crystal structure of *A. fumigatus* FKBP12 (purple) bound to FK520 (gray) characterized at a resolution of 1.7 Å. (C) Higher resolution focus of FK520 bound to the FKBP12 ligand-binding pocket. Both the C22 and the C21 of FK520 approach key fungal-specific residue F88 in the 80s loop of FKBP12.

Due to inherent challenges in forming the full-length ternary complex for crystallography, we next generated a novel “mini-calcineurin” as an alternative approach to capture FK520 bound to human FKBP12 and calcineurin **(Supplementary Fig. 1)**. The mini-calcineurin construct consists of the full length CnB subunit fused C-terminally to the calcineurin B-binding helix (BBH, *Af* residues 335-370, *Hs* residues 337-372) of CnA, lacking the CnA catalytic domain present in previous structures. Constructs for both human and *A. fumigatus* were generated and high-resolution ternary structures of each protein bound to FK520 (PDB: 7U0T) and FK506 (PDB: 7U0U), respectively were characterized **(Table S1)**. Remarkably, overlays with bovine FKBP12-FK506-calcineurin (PDB: 1TCO) reveal consistent conformation of the mini-calcineurin structure demonstrating that the key molecular interactions between FKBP12-FK520 and calcineurin are largely attributable to the hydrophobic interface presented by CnB bound to the extended CnA alpha-helical arm. Root-mean-square deviation (RMSD) values for the overlay of the *A. fumigatus* mini-calcineurin was 0.90 Å, consistent with previously reported values of structural similarity(34). Human mini-calcineurin RMSD value was 0.57 Å, indicative of a high degree of structural similarity and that the truncation of CnA did not significantly alter the overall binding structure of the complex. Each of these novel FK520-bound structures represent a key addition to the tools necessary for structure-guided design of both FK506/FK520 analogs that are increased for fungal selectivity.

**Table 1.** Minimum Inhibitory/Effective Concentrations (MIC/MECs) for FK506/FK520 analogs.

|  | Organism tested for MIC/MEC (µg/mL) |  |  |
| --- | --- | --- | --- |
|  | *Cryptococcus neoformans* (H99)† | *Aspergillus fumigatus* (Af293) | *Candida albicans* (SC5314)†† |
| FK506 | 0.05 | 0.016 | 0.02 |
| FK520 | 0.05 | -- | 0.02 |
| JH-FK-01 | 1 | 1 | 0.4 |
| JH-FK-02 | 2 | 2 | 1.6 |
| JH-FK-03 | 1 | 1 | 0.8 |
| JH-FK-04 | 2 | 2 | 6.25 |
| JH-FK-05 | 1 | 2 | 0.8 |
| JH-FK-07 | 0.2 | 1.25 | 0.2 |

† *C. neoformans* MICs performed at 37 °C
†† *C. albicans* MICs performed in the presence of 1 µg/mL fluconazole
-- not tested

### Chemical synthesis of novel FK506 and FK520 analogs modified at the C22 position

Our previous study demonstrated that C22 modifications of FK506 can introduce fungal selectivity(34). To leverage the protein crystal structures of FKBP12 bound to both FK506 and FK520, we developed a one-step synthetic protocol for C22 modification compatible with either starting material (**Fig. 2A**). The C22 position of FK506 and FK520 can be selectively targeted through an easily scalable and high-yielding condensation reaction with an acylhydrazone under reflux conditions. Inspired by the success of the first-generation FK506 analog APX879, we re-synthesized this compound (JH-FK-01) and generated five additional C22-modified compounds (**Fig. 2B**). Of these five second-generation analogs, three were derived from FK506 (JH-FK-02, -03, -04) and two were derived from FK520 (JH-FK-05, -07). Each analog was validated with high resolution mass spectrometry (HRMS) and ^1^H NMR **(Table S2, Supplementary Fig. 4)**.

**Figure 2.**
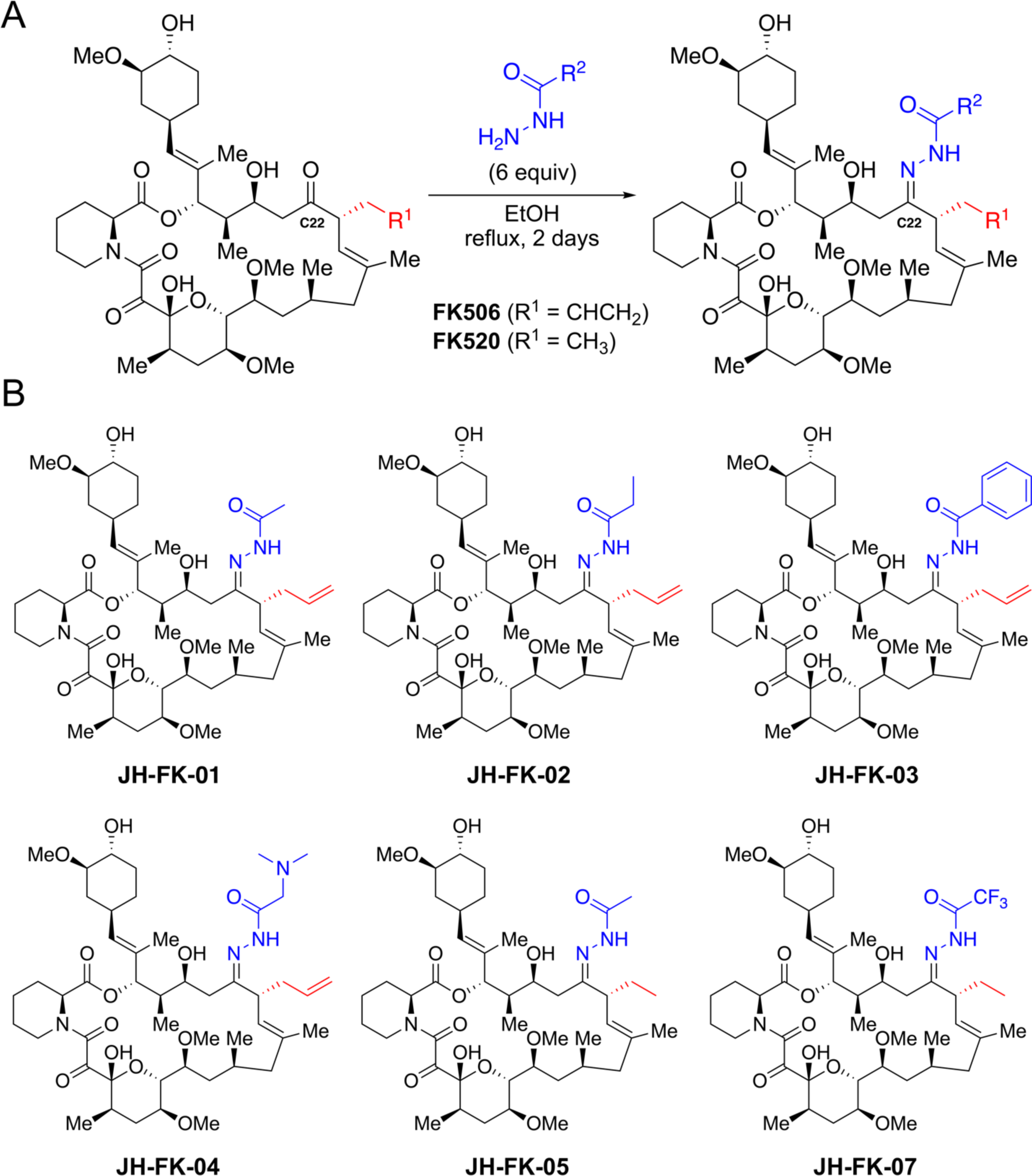
Chemical synthesis of novel FK506 and FK520 analogs. (A) One-step synthesis for C22-modifed analogs from FK506 or FK520 as starting material. (B) Panel of six FK506/FK520 analogs with acylhydrazone groups at C22. Acylhydrazones differ in R^2^ group. Compounds JH-FK-01, JH-FK-02, JH-FK-03, JH-FK-04 are derivatives of FK506. Compounds JH-FK-05 and JH-FK-07 are derivatives of FK520.

### Second-generation FK506 and FK520 analogs display broad-spectrum antifungal activity

Minimum inhibitory/effective concentrations (MIC/MECs) were determined for each of the novel analogs against the three major human fungal pathogens, *A. fumigatus*, *C. neoformans*, and *C. albicans*. Although all six analogs were reduced for antifungal activity compared to FK506, they were each effective at inhibiting growth in microbroth dilution assays **(Table 1)**. Along with their robust immunosuppressive activity, FK506 and FK520 exhibit remarkably potent antifungal activity in the range of 0.016-0.05 µg/mL which is far below the single-digit µg/mL MIC threshold typically necessary to translate to *in vivo* antifungal efficacy. Thus, analogs of FK506 and FK520 can tolerate up to 100-fold reduction in antifungal activity and still be considered for *in vivo* testing. For *C. neoformans*, MICs were the most uniform and ranged from 0.2 to 2 µg/mL which is between 4- to 40-fold reduced for antifungal activity. *C. albicans* MICs ranged from 0.2 to 6.25 µg/mL which is between 10- to 312.5-fold reduced for activity. Despite being one of the most difficult to treat fungal infections, all six analogs were active against *A. fumigatus* with MECs ranging from 1 to 2 µg/mL or a 62.5- to 125-fold reduction in activity.

Two FK520 analogs, JH-FK-05 and JH-FK-07, maintained high broad spectrum antifungal activity across the three species and JH-FK-07 had the lowest MICs for both *C. neoformans* and *C. albicans*. To assess the antifungal activity against other human fungal pathogens, we selected JH-FK-05 to test against a panel of pathogenic molds and drug-resistant *A. fumigatus* strains **(Table S3)**. JH-FK-05 registered MECs as low as 1 µg/mL for some of these highly recalcitrant species such as *Mucor circinelloides*, *Rhizopus oryzae,* and pan azole-resistant *A. fumigatus* (F14946).

To confirm that JH-FK-05 was fungal-FKBP12-dependent, we utilized a humanized *A. fumigatus* strain (*akuB^Ku80^* hFKBP12) expressing a codon-optimized human *FKBP12* gene from the *Affkbp12* native locus. We have previously shown that this hFKBP12-expressing strain is completely resistant to FK506 and that the activity of FK506 in *Aspergillus* is dependent on the presence of an FKBP12 that can form the FKBP12-inhibitor-calcineurin complex(34). Similar to FK506, JH-FK-05 has no detectable antifungal activity against the hFKBP12-expressing *A. fumigatus* strain suggesting that JH-FK-05 is working via the same mechanism of action **(Table S3)**.

### FK506 and FK520 analogs bind FKBP12 and form ternary inhibitory complex with calcineurin

To better understand the relationship between structure and activity of these analogs, we tested one FK506 analog, JH-FK-02, and one FK520 analog, JH-FK-05, for their binding dynamics to CnA and CnB by Biolayer Interferometry (BLI). Purified and biotinylated (Avi-tagged) FKBP12 from *A. fumigatus* and *C. albicans* were loaded onto streptavidin sensors in the presence of inhibitor, followed by titration of the respective CnA-CnB-calmodulin complex also in the presence of inhibitor. Data were double referenced using parallel reference sensors and sample blanks. Steady state and kinetic analyses confirm that the target complexes form in the presence of the analogs with similar affinities as the parent compounds, suggesting that the *in vivo* observations are due to inhibitory effects of the compound-mediated calcineurin-FKBP12 complexes **(Supplementary Fig. 2).**

### Some second-generation FK506 and FK520 analogs are increased for fungal specificity

The potent immunosuppressive activity of FK506 and FK520 is what precludes these highly antifungal compounds from being utilized clinically to treat invasive fungal infections. Therefore, it is essential to understand the level of immunosuppressive activity for each of these analogs to determine if these C22 modifications have improved upon the fungal selectivity. Utilizing a primary murine T cell model, we assessed IL-2 expression following growth in the presence of calcineurin inhibitors (**Fig. 3A**). Naïve, primary CD4+ T cells were collected from mice and grown for 72 hours in the presence of FK506, FK520, or an analog. Following stimulation with PMA and ionomycin, the proportion of cells producing IL-2, a key calcineurin-dependent cytokine, was measured with flow cytometry. All compounds generated a dose-dependent reduction in IL-2 production. FK506 and FK520 demonstrated the highest level of IL-2 inhibition and the effect of FK520 on IL-2 production was slightly reduced compared to that of FK506. JH-FK-02 and JH-FK-05 were the least immunosuppressive second-generation analogs with IC_50_ values of 42.6 nM (>380-fold increase compared to FK506) and 20.0 nM (>180-fold increase compared to FK506), respectively (**Fig. 3A**).

**Figure 3.**
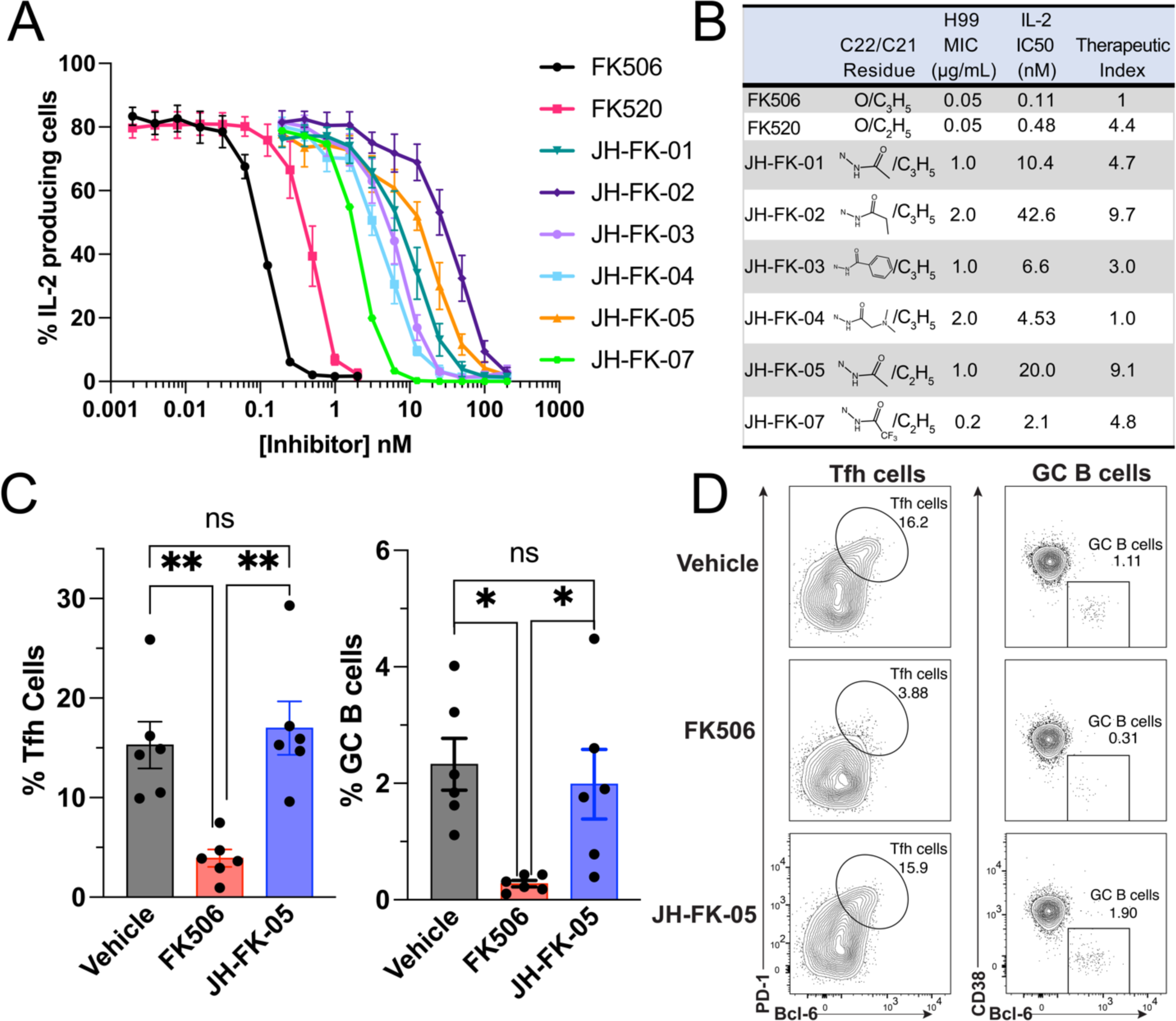
JH-FK-05 is reduced for immunosuppressive activity *in vitro* and *in vivo* and demonstrates fungal specificity. (A) *In vitro* immunosuppressive activity of calcineurin inhibitors was measured in a primary murine T cell model. Dose response curves were generated from IL-2 expression in cells exposed to increasing concentrations of inhibitors. All analogs were reduced for immunosuppression compared to controls FK506 (black) and FK520 (pink). Error bars indicate SEM. (B) Therapeutic Index scores for FK506/FK520 analogs were calculated by comparing the ratio of antifungal (MIC for *C. neoformans*) to immunosuppressive activity (IL-2 IC_50_) to that of FK506. FK506 therapeutic score was set to 1.0 and increasing scores indicate increasing fungal specificity. (C) *In vivo* immunosuppression of FK506 and JH-FK-05 was measured in a murine model of a T cell-dependent response. Animals were treated daily with vehicle, FK506 (5 mg/kg), or JH-FK-05 (60 mg/kg) beginning on day -1 via IP injection. NP-OVA was administered subcutaneously on day 0 to promote T cell-dependent proliferation of T follicular helper cells (Tfh) and germinal center B cells (GC B). Proportion of proliferated Tfh cells and GC B cells was measured in animal lymph nodes on day 7. Tfh and GC B cells are measured from TCRβ+ CD4+ CD44hi cells and TCRβ-CD19+ cells, respectively. (D) Representative flow cytometry plots are presented from individual animals in each treatment group. Gating indicates region containing Tfh cells (left, oval) or GC B cells (right, square). Error bars indicate standard deviation. For all plots, ** P<0.01, * P<0.05.

To assess the therapeutic potential for these compounds it was important to develop an approach to score and rank them based on antifungal and immunosuppressive activities. The Therapeutic Index score (TIscore) is a ratio of the immunosuppressive fold change to the antifungal fold change with all values relative to the baseline FK506 activity (**Fig. 3B**). A TIscore > 1.0 indicates an increase in fungal specificity because the fold change of the immunosuppressive reduction is greater than the antifungal activity reduction. As an example, JH-FK-05 is 20-fold reduced for antifungal activity but about 180-fold reduced for immunosuppression. Therefore, the TIscore of JH-FK-05 is 9.1 (TIscore=182/20). The top two analogs were JH-FK-02 (TIscore= 9.7) and JH-FK-05 (TIscore= 9.1). Due to the similar TIscore, JH-FK-05 was prioritized to advance to animal trials on the merit of more pronounced antifungal activity compared to JH-FK-02.

### JH-FK-05 is non-immunosuppressive in a murine model of T cell-dependent response

Based on the *in vitro* evidence of reduced immunosuppressive activity of JH-FK-05, we hypothesized that this reduction would translate into reduced immunosuppression in an *in vivo* model. Our murine model for immunosuppression measures the T cell-dependent proliferation of two immune cell subtypes in response to immunization with an antigen. Animals were treated daily via intraperitoneal (IP) injection with vehicle, 5 mg/kg FK506, or 60 mg/kg JH-FK-05 beginning on day -1 and ending on day 7. JH-FK-05 treatment was tolerated at a dose as high as 60 mg/kg and the final dose of JH-FK-05 was selected based on the maximum compound solubility (data not shown). On day 0, animals were immunized with NP-OVA to stimulate T cell-dependent signaling that would result in proliferation of T follicular helper (Tfh) cells and germinal center (GC) B cells. Animals were sacrificed on day 7 and the proportion of Tfh and GC B cells in the lymph nodes was measured. Treatment with 5 mg/kg of FK506 significantly reduced the proliferation of both Tfh and GC B cells (**Fig. 3B**). However, despite the much larger daily dose of 60 mg/kg, JH-FK-05 treatment did not reduce the presence of either cell type and was statistically indistinguishable from the vehicle treated animals. Representative flow cytometry plots capture the lack of both immune cell subtypes only in the FK506-treated animals (**Fig. 3D**). In addition to its lack of immunosuppressive activity *in vivo*, JH-FK-05 was well-tolerated by animals throughout the course of treatment.

### JH-FK-05 reduces pulmonary and brain fungal burden in models of cryptococcosis

The high *in vitro* antifungal activity of JH-FK-05 presented promising evidence that it might effectively reduce fungal burden in the context of murine infection. We employed two models of *C. neoformans* infection to test this hypothesis (**Fig. 4A**). First, in the intranasal model, animals were anesthetized and infected with 10^5^ cells of wild type H99 *C. neoformans* via instillation in the animal’s nares. These animals were monitored for 14 days and then sacrificed. This model tests the efficacy of JH-FK-05 to treat a pulmonary infection, which is the most common manifestation of early-stage cryptococcosis in patients. Second, the intravenous model approximates the disseminated stage of disease. Animals were infected with 10^4^ cells via injection into the lateral tail vein. Due to the advanced nature of the model and the rapidly disseminated disease, animals were sacrificed at 7 days and found to have a substantial CNS fungal burden. In both models, animals were treated once daily via IP injection beginning 3 hours after infection with either vehicle, 60 mg/kg JH-FK-05, 12 mg/kg fluconazole, or a combination of the two monotherapies. Fluconazole is a widely used antifungal drug targeting ergosterol biosynthesis for cell membrane integrity. Previous work has shown that calcineurin inhibitors are synergistic with fluconazole (37, 38). Additionally, the first generation FK506 analog, APX879, benefits from this synergistic interaction when analyzed *in vivo* in combination therapy with fluconazole in the pulmonary cryptococcosis model(34).

**Figure 4.**
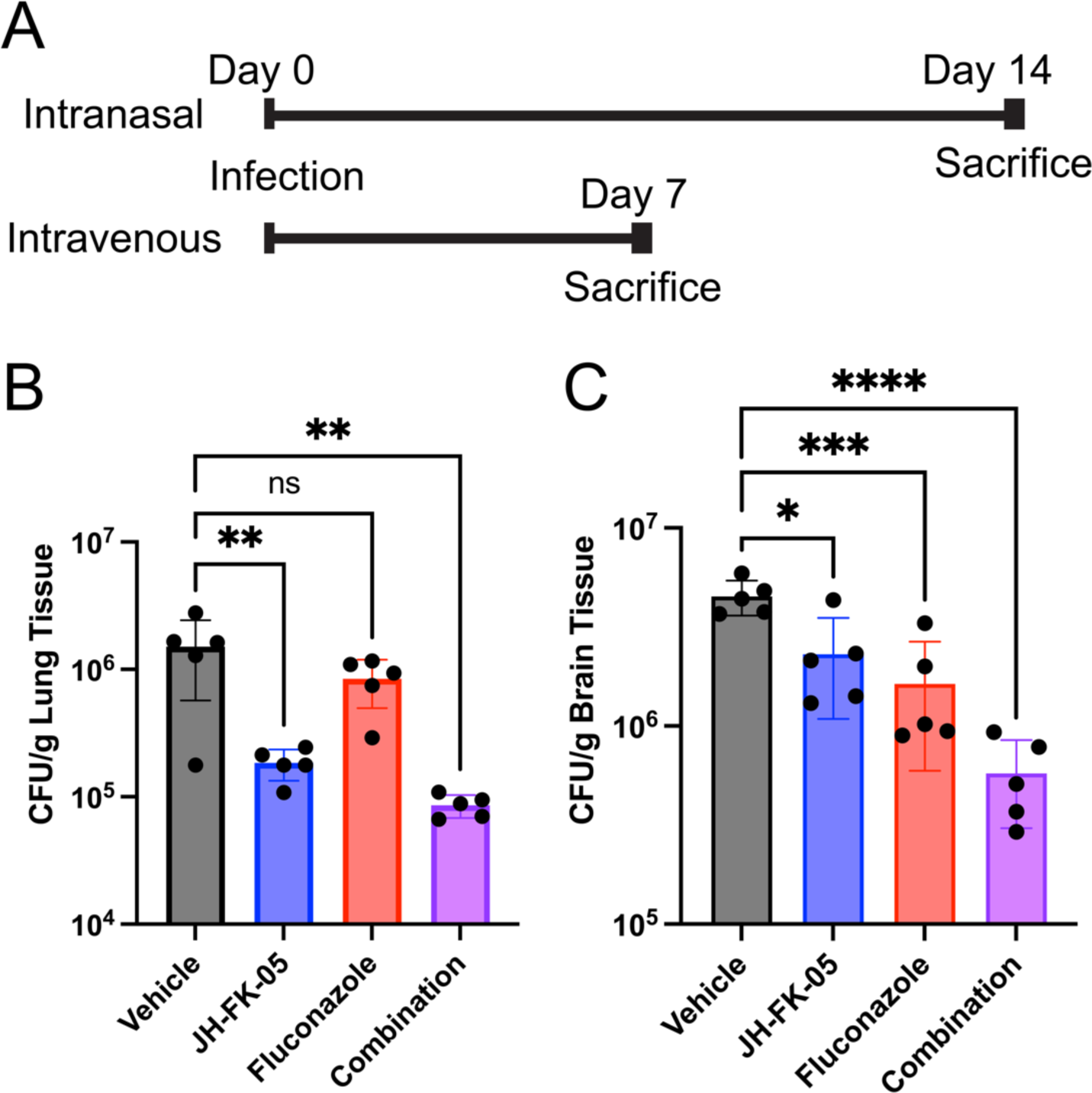
JH-FK-05 treatment reduces organ fungal burden in multiple models of *C. neoformans* infection. (A) Experimental timelines for intranasal and intravenous models of *C. neoformans* infection. For both models, treatment groups included vehicle (gray), JH-FK-05 60 mg/kg (blue), fluconazole 12 mg/kg (red), and combination (purple). Treatment began on day 0 via IP injection and continued daily until the indicated endpoint. Each treatment group contained 5 female A/J mice. (B) Lung tissue fungal burdens from animals in intranasal model. (C) Brain tissue fungal burdens from animals in intravenous model. For all plots, **** P<0.0001, *** P<0.001, ** P<0.01, * P<0.05. Error bars indicate standard deviation.

Like the immunosuppressive murine model, animals tolerated JH-FK-05 treatment for the full treatment period. Analysis of lung tissue collected from animals at day 14 showed that JH-FK-05 treatment alone reduced fungal burden by nearly 10-fold compared to vehicle-treated animals (**Fig. 4B**). The sub-therapeutic dose of fluconazole did not reduce the fungal burden alone but combination therapy with JH-FK-05 significantly reduced the lung fungal burden by over 10-fold. To assess if JH-FK-05 treatment could display therapeutic levels of activity after the infection had fully disseminated, the intravenous model was employed. A modest but significant 2-fold reduction (P<0.05) in brain fungal burden was detected in animals treated with JH-FK-05 alone (**Fig. 4C**). However, in combination, JH-FK-05 and fluconazole treatment results in a nearly 10-fold reduction in fungal burden after only 7 days of treatment. This demonstrates that not only can JH-FK-05 treat an aggressive pulmonary-stage infection, but it can cross the blood-brain barrier to directly treat a disseminated CNS infection. This is in accord with previous studies and known pharmacodynamic properties of FK506, which is known to cross the blood brain barrier(17). Together, these results demonstrate that JH-FK-05 is effective at reducing fungal burden in multiple stages of *C. neoformans* infection both as a monotherapy and in combination with the FDA-approved antifungal drug fluconazole.

### Treatment with JH-FK-05 in murine *C. neoformans* infection extends survival

To verify if this reduction seen in the fungal burdens of JH-FK-05 and combination-treated animals could translate into a meaningful therapeutic effect, the survival of animals infected via the intranasal route was assessed (**Fig. 5A**). As in the fungal burden model, these animals received daily treatment for only 14 days. Following the termination of treatment, animals were monitored for signs of decline, lethargy, and ultimately, moribundity. Untreated animals in this model typically survive around 3 weeks and similar results were observed here (vehicle median survival, 22 days). Corroborating the results of the fungal burden study, a significant extension of survival for both JH-FK-05 (median survival, 26 days) and fluconazole (median survival, 25 days) was observed. As with the fungal burden analysis, combination of JH-FK-05 and fluconazole had a potent therapeutic effect and extended survival of these animals to a maximum of 34 days (median survival, 31 days). Both monotherapy survival curves were significantly different from the combination therapy suggesting an additive or synergistic effect *in vivo* (**Fig. 5B**).

**Figure 5.**
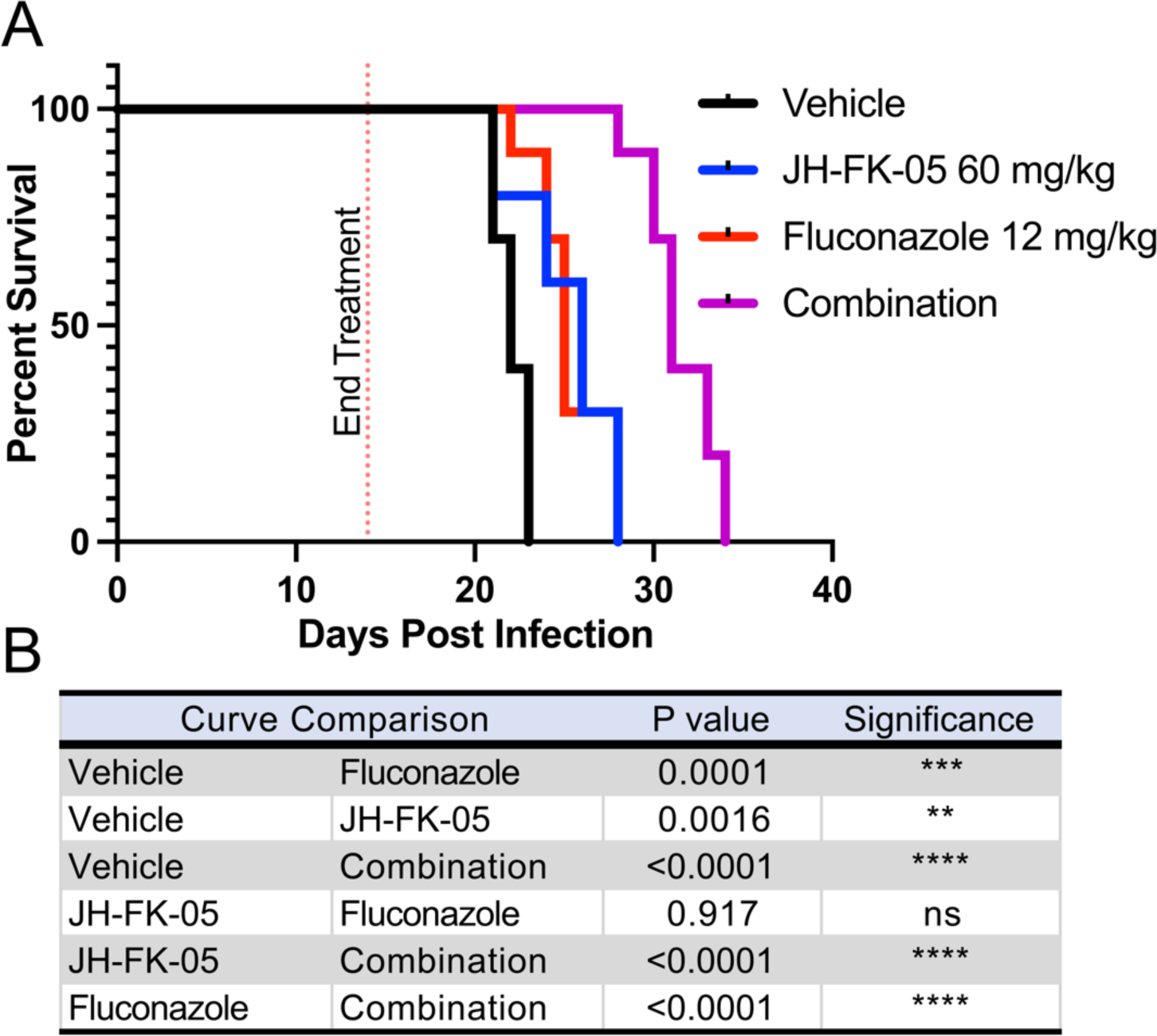
JH-FK-05 treatment extends animal survival in murine model of cryptococcosis. (A) Survival plot of animals infected with *C. neoformans* H99 through intranasal instillation. Treatment groups included vehicle (black), JH-FK-05 60 mg/kg (blue), fluconazole 12 mg/kg (red), and combination (purple). Animals received treatment via IP injection daily for 14 days and animals were monitored for health following treatment termination. Each treatment group consisted of 10 A/J female mice. (B) Survival curve pairwise comparisons by Log-rank (Mantel-Cox) test.

### Molecular dynamic simulations identify novel JH-FK-05 residues for future medicinal chemistry

Although JH-FK-05 has demonstrated robust *in vitro* and *in vivo* antifungal activity, additional improvements can be made to elevate the TIscore of this compound. To address this, molecular dynamic (MD) simulations of JH-FK-05 physical interactions with human vs fungal FKBP12 were conducted to identify contact residues that are more important for either species (**Fig. 6**)(36). A series of 500 nanosecond MD simulations between hFKBP12 and JH-FK-05 or *C. neoformans* FKBP12 and JH-FK-05 were simulated and differential contacts between JH-FK-05 and the two FKBP12 proteins were observed. Simulation stability was monitored with alpha carbon RMSD, radius of gyration, and center of mass distance **(Supplementary Fig. 3)**. Both the significant protein residues and the JH-FK-05 molecular residues that were identified are labeled on the respective structure (**Fig. 6**, **Table S4)**. There were 4 residues of JH-FK-05 that were more significant for *C. neoformans* contact (**Fig. 6**, cyan label). Interestingly, there were only 2 residues that were more significant for the JH-FK-05 and hFKBP12 interaction (**Fig. 6**, magenta labels). These residues are both in the C2-N7 ring of the molecule and represent potential future target areas for medicinal chemistry to introduce further fungal specificity to JH-FK-05 and design a new generation of analogs.

**Figure 6.**
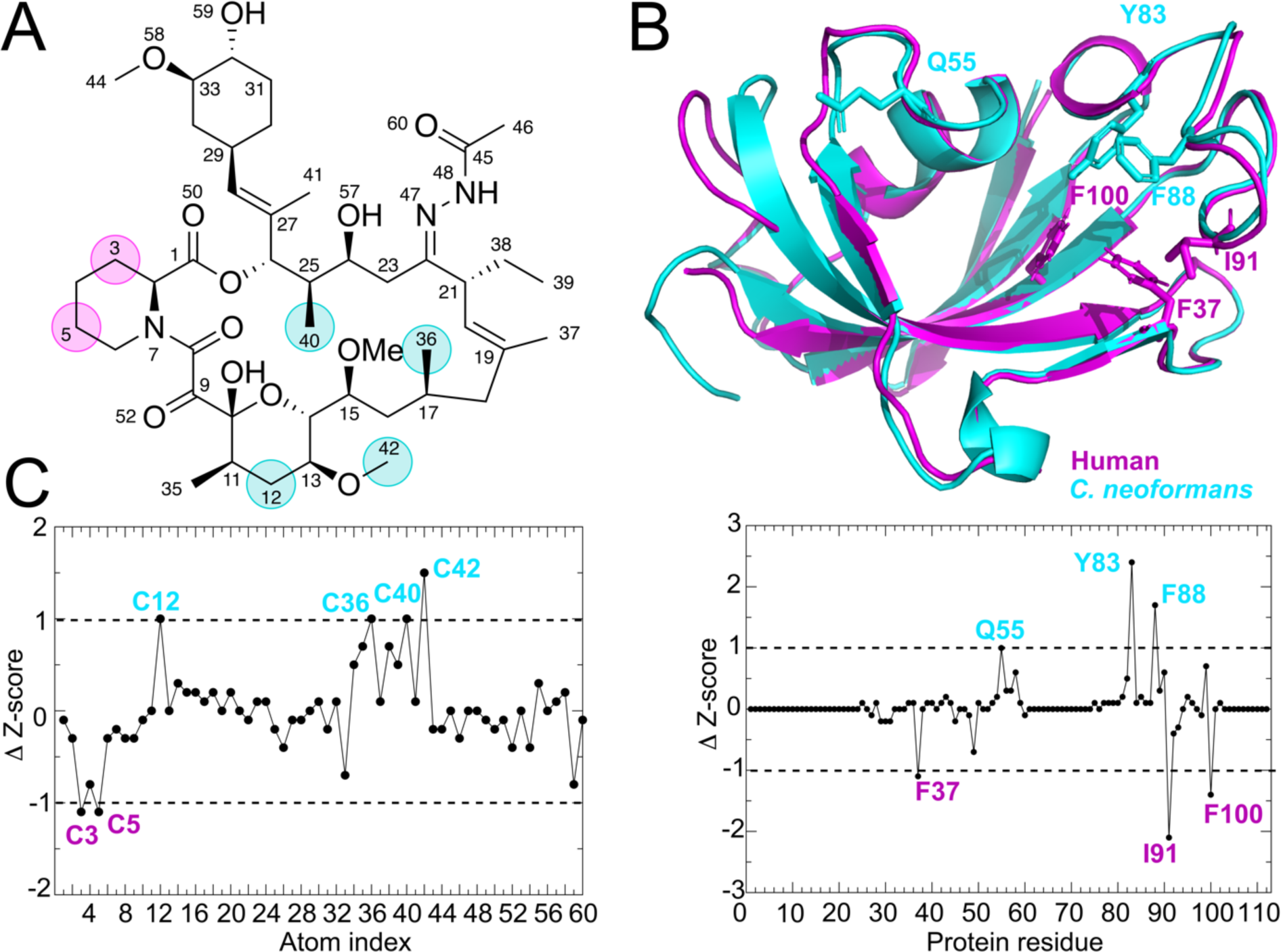
Ligand atoms and protein residue contacts observed in MD simulations. Analysis of the significance of the observed contacts of *h*FKBP12 and *C. neoformans* FKBP12 with JH-FK-05. (A) Stick representation of JH-FK-05 labeled for atoms making more significant contacts to *h*FKBP12 (magenta circles) and *C. neoformans* (cyan circles). (B) Residues of *h*FKBP12 (magenta) and *C. neoformans* (cyan) making more significant interactions to the ligand are shown in stick format on the ribbon protein structures. (C) Z-score graphs: black circles represent Z-scores of *C. neoformans* FKBP12 contacts minus *h*FKBP12 contacts. More significant contacts for *C. neoformans* (Δ Z-score >1) are labeled in cyan while contacts more significant for the human protein (Δ Z-score <1) are labeled in magenta.

## Discussion

In this study, a panel of second-generation FK506 and FK520 analogs were screened and JH-FK-05 was identified as a promising antifungal candidate that targets fungal calcineurin with a high degree of specificity. We also report the first fungal FKBP12-FK520 protein crystal structure at a high resolution of 1.7Å. Much of this work was made possible by the structure-guided design and development of a first generation FK506 analog, APX879(34). Crystal structures of fungal FKBP12 proteins revealed a key conserved difference in the 80s loop that may promote fungal selectivity in molecules modified at the C22 position of FK506. The *A. fumigatus* FKBP12-FK520 structure clearly shows that both the C22 and C21 residues of FK520 approach the Phe88 residue (**Fig. 1C**). This proximity and potential for a steric clash suggested that modifications at the C22 position on both FK506 and FK520 could introduce a high degree of selectivity.

Establishing the therapeutic window for these calcineurin inhibitors was an essential component of developing small molecules that were fungal-specific. Because FK506 and FK520 already have exceedingly high antifungal activity in addition to their immunosuppressive activity, the goal was to reduce the immunosuppression while preserving antifungal activity to shift the balance in favor of this therapeutic window. Both inhibitors have stronger antifungal activity against *C. neoformans* than the front-line treatments for cryptococcosis, amphotericin B (MIC 0.125 µg/mL) and fluconazole (3 µg/mL) **(Table 1)** (39, 40). Due to the elevated starting point for both of these compounds, we were able to afford a significant reduction in the antifungal activity seen in each of our analogs. Two compounds were identified, JH-FK-02 and JH-FK-05, that demonstrate a nearly 10-fold increase in Therapeutic Index score overall and are only 40- and 20-fold reduced for antifungal activity, respectively (**Fig. 3B**). These compounds also helped establish a “threshold” for prioritizing future compounds for advancement into animal models. Based on the high antifungal activity of FK506/FK520 and the reported MIC/MECs for clinically utilized antifungal drugs, we estimate that our analogs can tolerate up to a 100-fold reduction in antifungal activity and still perform well in infection models (39, 40).

JH-FK-05 was selected for further *in vitro* mold testing and *in vivo* animal models due to its higher degree of antifungal activity compared to JH-FK-02 (**Fig. 3B**). Remarkably, despite a daily dose at 12 times that of FK506, there was no detectable change in the T cell-dependent response compared to vehicle-treated animals (**Fig. 3C**). This shift in the immunosuppressive activity facilitates the *in vivo* dosing of JH-FK-05 at a high level of 60 mg/kg and was limited primarily by solubility and not by tolerability. Comparatively, in the same murine infection models we have shown that a lower, 20 mg/kg dose of FK506 is lethal to animals(34). Although not compared directly in this study, JH-FK-05 represents a significant reduction in toxicity and improvement in efficacy to be within the therapeutic window. Pharmacokinetic investigation of JH-FK-05 treatment could lead to an optimization of the dosage frequency and total dose in the future. Additionally, histopathological analysis of infected tissue will determine the level of residual fungal growth and assess the recruitment of immune cells to the site of infection. Due to the high degree of selectivity observed with JH-FK-05, it is possible that treatment will not be limited by toxicity but instead be limited by the solubility and delivery route of the treatment. Fortunately, both FK506 and FK520 are orally active compounds in both humans and rodents(41). Additional investigation will determine if JH-FK-05 can be orally administered but the pharmacokinetic properties of the parent molecule FK520 suggests that this is likely to be the case. The development of an orally-active antifungal compound with broad-spectrum activity would be a major addition to the currently limited antifungal armamentarium(4).

We utilized two different models of *Cryptococcus* infection to address the key stages of infection experienced by patients. The intranasal infection model is similar to the progression of disease observed in most patients that begin with a pulmonary infection. The intravenous model allowed us to test if JH-FK-05 can treat an already disseminated infection and exert activity across the blood-brain barrier (BBB) (**Fig. 4**). FK506 is known to cross the BBB so it was validating to observe JH-FK-05 reduce fungal burden in the brain of IV-infected mice(17). CNS engagement and antifungal activity across the BBB are critical factors in developing treatments with therapeutic potential for the treatment of cryptococcosis. There is an increasing need for treatments that can directly address the CNS stage of infection(4). Without early intervention, the majority of pulmonary *C. neoformans* infections will disseminate to become aggressive CNS infections. In these cases, high rates of mortality in all groups of patients diagnosed with cryptococcal meningitis is a direct result of treatments failing to act on the disseminated versus the localized pulmonary disease stage (42, 43). JH-FK-05 represents a promising opportunity to address both early and late-stage *C. neoformans* infections.

In each of our infection models we demonstrated that combination treatment with fluconazole increased the potency of JH-FK-05 (**Fig. 4** and **Fig. 5**). In fact, the gold standard of treatment for *C. neoformans* infections is a combination therapy of amphotericin B and 5-fluorocytosine(44). Other antifungal combination therapies that have shown efficacy in the clinic include fluconazole+amphotericin B (candidemia) and caspofungin+amphotericin B (mucormycosis)(45). Due to rising rates of antifungal drug resistance, combination therapies that generate additive or synergistic interactions are essential in treatment plans. We suggest that the efficacy of JH-FK-05 as both a monotherapy and a combination therapy with a clinical antifungal drug is an exciting development and a promising indicator that this approach could lead to a treatment with substantial clinical impact.

Multiple groups have developed biosynthetic analogs of FK506 by deleting genes in the FK506 biosynthetic cluster of the *Streptomyces* organism(46). After the macrolide base of FK506 is produced by polyketide synthase (PKS) steps, several enzymes introduce post-PKS modifications to the macrolide ring. Deletion of genes encoding these enzymes (*fkbM*, *fkbD*) resulted in different FK506 analogs with varying degrees of immunosuppressive and antifungal activity(38). The scalability of this approach is extremely promising. Although this strategy is ultimately limited in rational design, future combinatorial synthetic approaches can unlock the medicinal chemistry at sites across the FK506 molecule. Most biosynthetic analogs are compatible with our procedure to modify the C22 position of FK506/FK520 and could be utilized as starting material to synthesize a new generation of analogs through a combinatorial biosynthetic-medicinal chemistry strategy.

Future second-generation FK520 analog development will also be informed by MD simulations that identify target residues on the molecule that are more important for fungal versus mammalian interaction. Data from MD simulations performed with JH-FK-05 binding human versus *C. neoformans* FKBP12 revealed multiple residues that were increased for fungal specificity (**Fig. 6**). This approach has successfully identified areas of the first-generation FK506 analog, APX879, that are enriched for contact with only the human FKBP12(36). Modifications made to JH-FK-05 could further increase its TIscore and drive the therapeutic potential even further. This approach can be applied to any additional analogs that are developed in the future as potential starting material for combinatorial synthetic strategies at both C22 and C21 positions. Our pre-clinical success with JH-FK-05 indicates that iterative and structure-informed design of FK506 analogs is a promising strategy to develop fungal-specific calcineurin inhibitors as therapeutics.

## Materials and Methods

### Protein Production

See Appendix for full methods on protein production and purification for X-ray crystallography. *A. fumigatus* FKBP12 P90G was expressed and purified as previously described(34).

### Complex of *Af*CnA-CnB + *Af*FKBP12-P90G with FK506 and *Hs*CnA-CnB + Bio-*Hs*FKBP12 with FK520

Complexes were formed by first mixing FKBP12 with 1.5x FK506/FK520 at 4 °C for 30 min followed by addition of CnA-CnB. The combined sample was incubated for an additional 30 min at 4 °C then injected onto a Superdex 200 column (GE Healthcare) equilibrated in 25 mM Tris pH 8.0, 200 mM NaCl, 1 mM TCEP, and 5 mM CaCl2. Fractions containing the 1:1 complex were pooled and concentrated to 20-24 mg/mL.

### Structure determination by X-ray crystallography

*A. fumigatus* FKBP12 was pre-incubated at room temperature for 30 min with 1.6 mM FK520 (100 mM DMSO stock, compound to protein ratio of 1.26) prior to being placed in sparse matrix crystallization trials using sitting drop vapor diffusion at 18.2 mg/mL. Crystallization trays were stored at 14 °C and crystals grew over the course of 4 months. Crystals were grown in JCSG+ (Rigaku Reagents) condition A1 containing 0.2 M Lithium Sulfate, pH 4.5 and 50% v/v PEG400, which also acted as a cryo-protectant during harvest. The dataset was collected on 9/16/2021 at the synchrotron APS beamline 21-ID-F on a MAR 300 CCD X-ray detector. Two copies of the compound-bound FKBP12 were placed per asymmetric unit (PDB ID:7U0S). The structure was solved by molecular replacement using a previously solved structure of *A. fumigatus* FKBP12 bound to FK506 (PDB: 5HWC). Molecular graphics and analyses performed with UCSF Chimera, developed by the Resource for Biocomputing, Visualization, and Informatics at the University of California, San Francisco, with support from NIH P41-GM103311(47).

### Crystallization conditions of *Af*CnA-CnB + *Af*FKBP12-P90G with FK506 and *Hs*CnA-CnB + Bio-*Hs*FKBP12 with FK520

Proteins were placed in sparse matrix crystallization trials using sitting drop vapor diffusion. Crystallization trays were stored at 14 °C. Crystals of *Af*CnA-CnB + *Af*FKBP12-P90G with FK506 grew after 1 month in JCSG+ (Rigaku Reagents) condition C6 containing 0.1 M Sodium Phosphate dibasic/Citric acid) pH 4.2 and 40% v/v PEG300 (also cryo-protectant). Crystals of *Hs*CnA-CnB + Bio-*Hs*FKBP12 with FK520 grew after 1 day in MCSG-1 (Microlytics) condition C5 containing 0.2 M Magnesium Acetate and 20% w/v PEG3350 (Cryo-preserved in 20% ethylene glycol). Datasets were collected on 12/10/2020 at the synchrotron APS beamline 21-ID-F on a MAR 300 CCD X-ray detector. A single copy of each complex was placed per asymmetric unit (*Hs*CnA-CnB + Bio-*Hs*FKBP12 with FK520 PDB ID:7U0T; *Af*CnA-CnB + *Af*FKBP12-P90G with FK506 PDB ID:7U0U). The structure of *Hs*CnA-CnB + Bio-*Hs*FKBP12 with FK520 was solved by molecular replacement using a previously solved structure of *B. taurus* CnA, CnB and FKBP12 bound to FK506 (PDB: 1TCO), and the structure of *Af*CnA-CnB + *Af*FKBP12-P90G with FK506 was solved by molecular replacement using a previously solved structure of *A. fumigatus* CnA, CnB and FKBP12 bound to FK506 (PDB: 6TZ7).

Diffraction data were reduced and scaled with XDS/XSCALE(48). Structures were solved by molecular replacement using the program Phaser in the Phenix program suite(49). Structures were refined using iterative cycles of TLS and restrained refinement with Phenix Refine and model-building using COOT(50). The final structures were validated using Molprobity and deposited in the Protein Data Bank (51-53). Diffraction data and refinement statistics are listed in **Table S1**. Diffraction images are available on Integrated Resource for Reproducibility in Macromolecular Crystallography (http://proteindiffraction.org/) (54, 55).

### Synthesis of FK506 and FK520 analogs

FK506 and FK520 monohydrate starting materials were purchased from APIChem Technology (AC-32477 and AC-5267). FK506 and FK520 analogs were prepared as follows: To a solution of FK506 or FK520 in anhydrous EtOH (0.1 M) was added acylhydrazine (6.0 equiv) at 25 °C. The resulting mixture was refluxed under N2 atmosphere. After stirring for 36-48 h, the reaction mixture was cooled to 25 °C and concentrated *in vacuo*. The resulting mixture was diluted with EtOAc and H2O. The layers were separated, and the aqueous layer was extracted with EtOAc. The combined organic layers were washed with brine, dried over anhydrous Na2SO4, and concentrated *in vacuo*. The residue was purified by silica gel column chromatography. All compounds were confirmed by HRMS and ^1^H NMR **(Table S2, Supplementary Fig. 4)**.

### Antifungal susceptibility testing

*In vitro* antifungal activity of FK506, FK520, and analogs was assessed in RPMI 1640 (Sigma-Aldrich) for all strains tested. Minimum inhibitory/effective concentrations (MIC/MEC) for each drug were measured using modified Clinical and Laboratory Standards Institute (CLSI) M38-A2 and M27-A3 standard *in vitro* antifungal susceptibility protocols. Broth microdilution assays for MIC/MEC assessment were performed as previously described(34). All antifungal assays with *C. neoformans* were performed at 37 °C. Antifungal susceptibility testing for *C. albicans* was performed in the presence of a uniform 1 µg/mL concentration of fluconazole to sensitize the organism to calcineurin inhibition.

### Biolayer Interferometry (BLI) Binding Experiments

BLI experiments were completed using either *A. fumigatus* or *C. albicans* FKBP12 and CnA/CnB/calmodulin complexes, along with compounds FK506, APX879, FK520, JH-FK-02, and JH-FK-05. Initial buffer (25 mM Tris, 200 mM NaCl, 1 mM TCEP, pH 8.0) was prepared using 1 M Tris-HCl pH 8.0 (Teknova #T1080), 5 M Sodium Chloride (Teknova #S0251), TCEP (Soltec Ventures Inc #M115), and was readjusted to pH 8 before sterile filtering. BSA-containing buffer was prepared by addition of BSA stock (30% in DPBS, Millipore Sigma #A9576) into the initial buffer to a final concentration of 1%. Compound-containing assay buffers were prepared by addition of compound stock (100 mM in DMSO-*d*6) into BSA-containing buffer to a final concentration of 10 µM. FKBP12 solutions were prepared using a two-step process, where FKBP12 was first diluted to 100 µg/mL with BSA-containing buffer, followed by dilution to 10 µg/mL with compound-containing assay buffer. Initial 2500 nM CnA/CnB/calmodulin complexes were prepared by direct dilution of protein stock with compound-containing assay buffer. Serial 1:1 dilutions were then performed with compound-containing assay buffer to prepare the remaining samples in the series (full series: 2500 nM, 1250 nM, 625 nM, 312.5 nM, 156.25 nM, 78.125 nM, 39.0625 nM, and 0 nM [sample blank]). FKBP12 and CnA/CnB/calmodulin complexes were incubated overnight prior to experiments.

Experiments were performed on an Octet Red96e instrument (Sartorius) utilizing Octet Data Acquisition software version 11.1.1.19, equilibrated to 25 °C. Streptavidin biosensors (SA, Sartorius #18-5019) were hydrated for a minimum of 30 min in compound-containing assay buffer prior to the start of each experiment. Each experiment consisted of two identical assays, each performed using separate biosensors: one using sensors loaded with FKBP12 (sample sensors), and the other using unloaded sensors (parallel reference sensors). Each assay consisted of the following steps: initial baseline (120 s, compound-containing assay buffer); FKBP12 loading (170 s for *Aspergillus*, 180 s for *Candida*); sample baseline (120 s, compound-containing assay buffer); CnA/CnB/calmodulin complex association (300 s); dissociation (900 s, same compound-containing assay buffer wells as the sample baseline step); sample baseline/association/dissociation cycle 2 (same times and wells as previous steps); sample baseline/association/dissociation cycle 3 (same times and wells as previous steps). Data were acquired for FKBP12-loaded sensors first, parallel reference sensors second.

Data was processed and analyzed using ForteBio Data Analysis software version 11.1.3.10. Data was processed using the Double Reference subtraction option, with Y axis aligned to baseline (from 0.1 to 119.8 s), inter-step correction aligned to dissociation, and Savitzky-Golay filtering active. Kinetics data was analyzed using the Association and Dissociation Curve Fitting option utilizing a 1:1 model. Fitting was performed using the Global (Full) option with Rmax Unlinked by Sensor (the one exception being *Aspergillus* sample JH-FK-05, which wouldn’t process cleanly unless the Rmax Linked option was selected instead). The Window of Interest spanned the full association and dissociation steps of each experiment. Steady State Analysis was completed using the Response Average from 290.0 – 295.0 s of the association step (because some binding responses did not reach equilibrium).

### *In vitro* immunosuppressive activity testing

Spleen and lymph nodes were collected from C57BL/6 mice and homogenized through a 40 µm filter. Pan-CD4+ T cells were enriched using MagniSort Mouse CD4 T cell Kit (eBioscience) according to the manufacturer’s protocol. Thereafter, CD4+ CD25-CD44lo CD62Lhi naïve T cells were FACS-sorted. The naïve CD4+ T cells were grown in Iscove’s Modified Dulbecco’s Medium (IMDM) supplemented with glutamine, penicillin, streptomycin, gentamicin, 2-mercaptoethanol, and 10% FBS. Cells were then cultured on anti-hamster IgG coated plates in the presence of hamster anti-CD3 epsilon and anti-CD28 antibodies (BD Biosciences), neutralizing anti-IL-4 antibody (eBioscience), recombinant IL-12 (10 ng/mL), and recombinant IL-2 (50 U/mL) for 72 h. During this period, cells were grown in the presence of 2x serially diluted FK506, FK520, or analog suspended in DMSO for 72 h. During the last 4 h of culture, phorbol 12-myristate 13-acetate (PMA; Sigma), ionomycin (Sigma), and GolgiStop (BD Biosciences) were added to the culture to facilitate detection of intracellular cytokines via flow cytometry.

Live cells were stained with FITC-conjugated anti-CD4 antibody (eBioscience) and Fixable Viability Dye eFluor 506 (eBioscience) on ice, followed by fixation and permeabilization using the Foxp3 Transcription Factor Staining Kit (eBioscience). Intracellular staining was performed at room temperature using PE-conjugated anti-IL-2 antibody (eBioscience), followed by analysis with a BD LSRFortessa X-20 flow cytometer.

### Preparation of compounds for *in vivo* studies

For *in vivo* studies on immunosuppressive activity, FK506 was purchased from Duke Pharmacy as Prograf (tacrolimus, Astellas) and was used as a solution of 1 mg/mL. Vehicle was prepared as a sterile solution of 90% PBS and 5% Kolliphor EL (Sigma C5135) and 5% ethanol. JH-FK-05 stock solution of 120 mg/mL was prepared as a powder dissolved in sterile 50% Kolliphor EL and 50% Ethanol. For administration to animals, JH-FK-05 stock was diluted to 12 mg/mL and 10% Kolliphor/Ethanol in sterile PBS. Fluconazole was purchased from Duke University Hospital Pharmacy as a sterile solution of fluconazole 2 mg/mL in PBS.

### *In vivo* immunosuppressive activity assessment of JH-FK-05

Groups of 6 female C57BL/6 mice (Jackson Labs) were treated daily by intraperitoneal (IP) injection with about 100 µl vehicle, 5 mg/kg FK506, or 60 mg/kg JH-FK-05 beginning on day -1. On day 0 mice were immunized with NP-OVA in alum via subcutaneous injection. Mice were treated and monitored daily until they were sacrificed on day 7. Draining lymph nodes were harvested on day 7 and the populations of T follicular helper cells and GC B cells were analyzed by flow cytometry. Tfh and GC B cells are measured from TCRβ+ CD4+ CD44hi cells and TCRβ-CD19+ cells, respectively.

### *C. neoformans* cell preparation for murine infection

Wild type H99 *C. neoformans* cells were grown overnight at 30 °C in a roller drum in YPD media. Cells were pelleted at 3,000 rpm and washed 3 times with sterile PBS. H99 cells were counted with a hemacytometer and diluted to 2x10^6^ cells/mL in PBS for pulmonary model and 2x10^5^ cells/mL in PBS for disseminated model.

### Intranasal model of pulmonary cryptococcosis

Female A/J (3- to 4-week-old; Jackson Labs) mice were anesthetized utilizing an isoflurane chamber. While anesthetized, 50 µL of 2x10^6^ cells/mL suspension was dripped onto nares of mice, allowing them to inhale full inoculum. Treatment began 3 h following infection with IP dosing of vehicle, 60 mg/kg JH-FK-05, 12 mg/kg fluconazole, or combination. Animals were treated daily for 14 days. After 14 days, animals were either sacrificed for fungal burden analysis or monitored for health and survival until reaching a humane endpoint. Survival analysis was plotted and performed in GraphPad Prism 9. Statistical significance was determined using Log-rank (Mantel-Cox) tests.

### Intravenous model of disseminated cryptococcosis

Male CD1 mice (3- to 4-week-old; Charles River) were restrained with a 50 mL conical tube holes at both ends for ventilation. Animal tails were warmed under heat lamp to enhance vasodilation and sterilized with alcohol wipes. A 50 µL suspension of 2x10^5^ cells/mL was injected into the lateral tail vein with an insulin syringe. Treatment proceeded as described in intranasal infection model until day 7 when animals were sacrificed for fungal burden analysis.

#### *Cryptococcus* fungal burden analysis

Mice were euthanized by CO_2_ inhalation at the pre-determined experimental endpoint. Organs from each animal were harvested and homogenized in sterile PBS using steel beads (Qiagen) and a bead beater. Organ homogenate was serially diluted in PBS. 100 µL of each dilution was plated to antibiotic plates (YPD 50 µg/mL ampicillin, 30 µg/mL chloramphenicol), incubated 48 to 72 h at 30 °C, and assessed for colony forming units (CFUs). CFU per gram of organ was plotted using GraphPad Prism 9. Statistical significance was calculated using an ordinary one-way ANOVA with Tukey’s multiple comparisons test.

### Molecular Dynamic simulations

MD simulations were performed to provide a better representation of the protein’s conformational flexibility and to more accurately characterize the proteins’ solution structure bound to JH-FK-05. Crystal structures were used as the starting conformations: *h*FKBP12 bound to FK506 (PDB: 1FKF) and *cn*FKBP12 (PDB: 6TZ8 – with calcineurin A/B removed) (34, 56). JH-FK-05 small molecule parameter and topology files were downloaded and created utilizing the Automated Topology Builder (ATB) and repository (57, 58). All molecular dynamic (MD) simulations were performed with the GROMACS 5.0.1 software package utilizing 6 CPU cores and one Nvidia Tesla K80 GPU(59). The single starting conformations used for all the MD simulations were resulting X-ray-characterized crystal structures noted above. MD simulations were performed with the GROMOS54a7 force field and the flexible simple point-charge water model. The initial structures were immersed in a periodic water box with a dodecahedron shape that extended 1 nm beyond the protein in any dimension and were neutralized with counter ions. Energy minimization was accomplished through use of the steepest descent algorithm with a final maximum force below 100 kJ/mol/min (0.01-nm step size, cutoff of 1.2 nm for neighbor list, Coulomb interactions, and Van der Waals interactions). After energy minimization, the system was subjected to equilibration at 300 K and normal pressure for 1 ns. All bonds were constrained with the LINCS algorithm (cutoff of 1.2 nm for neighbor list, Coulomb interactions, and Van der Waals interactions). After temperature stabilization, pressure stabilization was obtained by utilizing the v-rescale thermostat to hold the temperature at 300 K and the Berendsen barostat was used to bring the system to 10^5^-Pa pressure. Production MD calculations (500 ns) were performed under the same conditions, except that the position restraints were removed, and the simulation was run for 500 ns (cutoff of 1.1, 0.9, and 0.9 nm for neighbor list, Coulomb interactions, and Van der Waals interactions). These MD simulations were repeated 6 times. Carbon alpha root mean square deviation (Cα-RMSD), radius of gyration (Rg), and center of mass (COM) all confirmed stability and accuracy of the MD simulations by stabilizing after a 100-ns equilibration period (in most cases) allowing for the analysis of the last 400 ns of the simulations. Only one of the 12 calculations (hFKBP12:JHFK05 complex MD sim #3) was rejected from the analysis due to unstable Cα-RMSD, Rg, and/or COM **(Supplementary Fig. 3)**. GROMACS built-in and homemade scripts were used to analyze the MD simulation results and averaged over the 6 simulations. All images were produced using PyMOL(60). Atom indices for JHFK05 is provided in **Table S4**.

## Acknowledgements

This project has been funded in whole or in part with grants from the National Institute of Allergy and Infectious Diseases, National Institutes of Health, Department of Health and Human Services, R01 AI112595-04, NIH predoctoral fellowship F31 AI150120, R56 AI112595-05, R01 GM115474, and HHSN272201700059C. The authors thank the many members of Duke University and UCB Biosciences for their contributions to the research and review of this manuscript.

## Supplementary Figure Legends

**Supplementary Figure 1.**
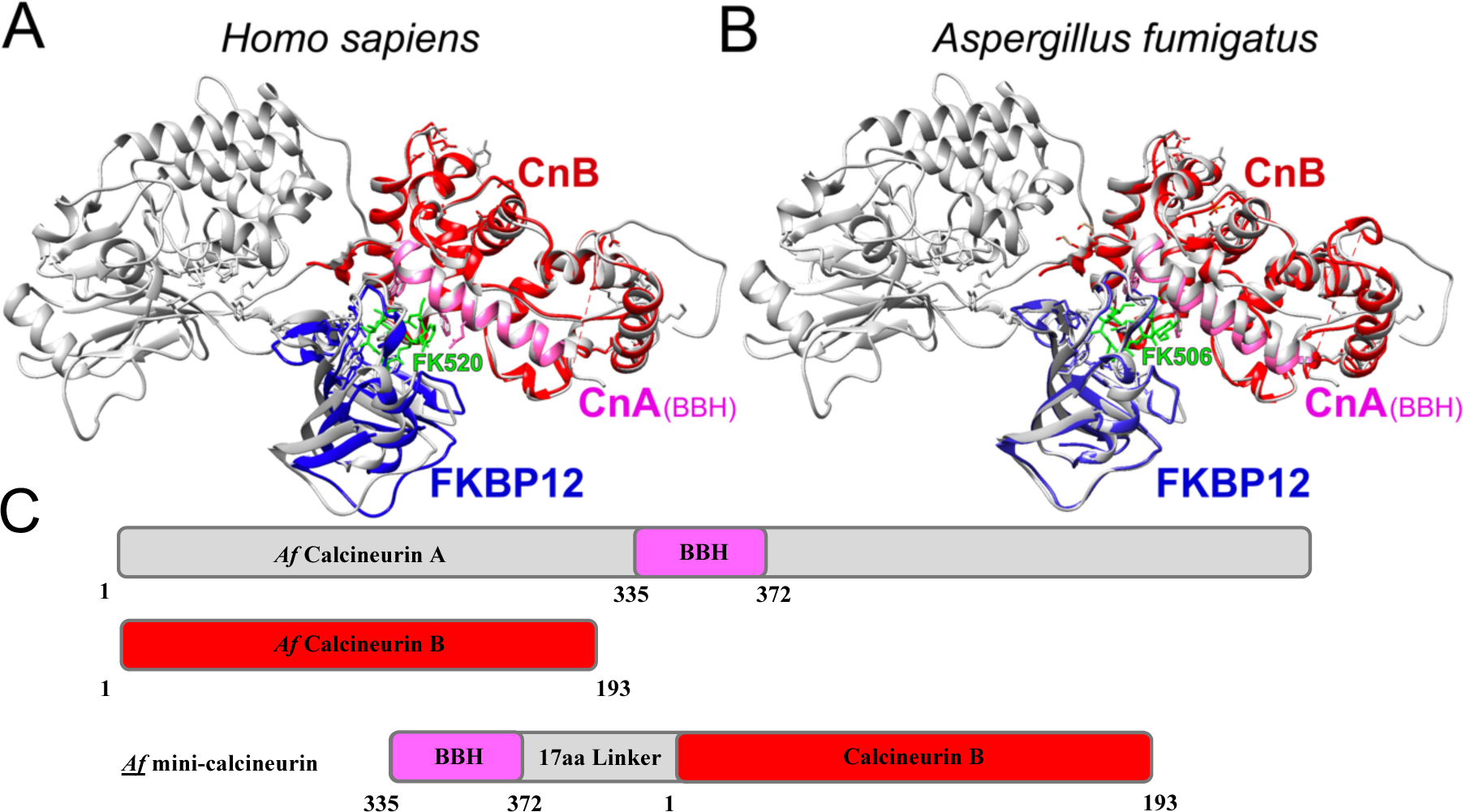
Mini-calcineurin crystal structures of human and *A. fumigatus* protein bound to FK520 and FK506. Protein crystal structures for each organism presented with overlay of bovine full length calcineurin ternary complex structure calcineurin-FKBP12-FK506 (PDB: 1TCO) in gray. (A) Structure of human mini-calcineurin complex (PDB: 7U0T) consisting of CnB (red) and CnA B Binding Helix (BBH, pink), FKBP12 (blue), and FK520 (green). (B) Structure of *A. fumigatus* mini-calcineurin complex (PDB: 7U0U) consisting of CnB (red) and CnA B Binding Helix (BBH, pink), FKBP12 (blue), and FK506 (green). (C) Protein expression construct of full length calcineurin subunits A and B and mini-calcineurin containing CnB and CnA BBH linked by 17 amino acids.

**Supplementary Figure 2.**
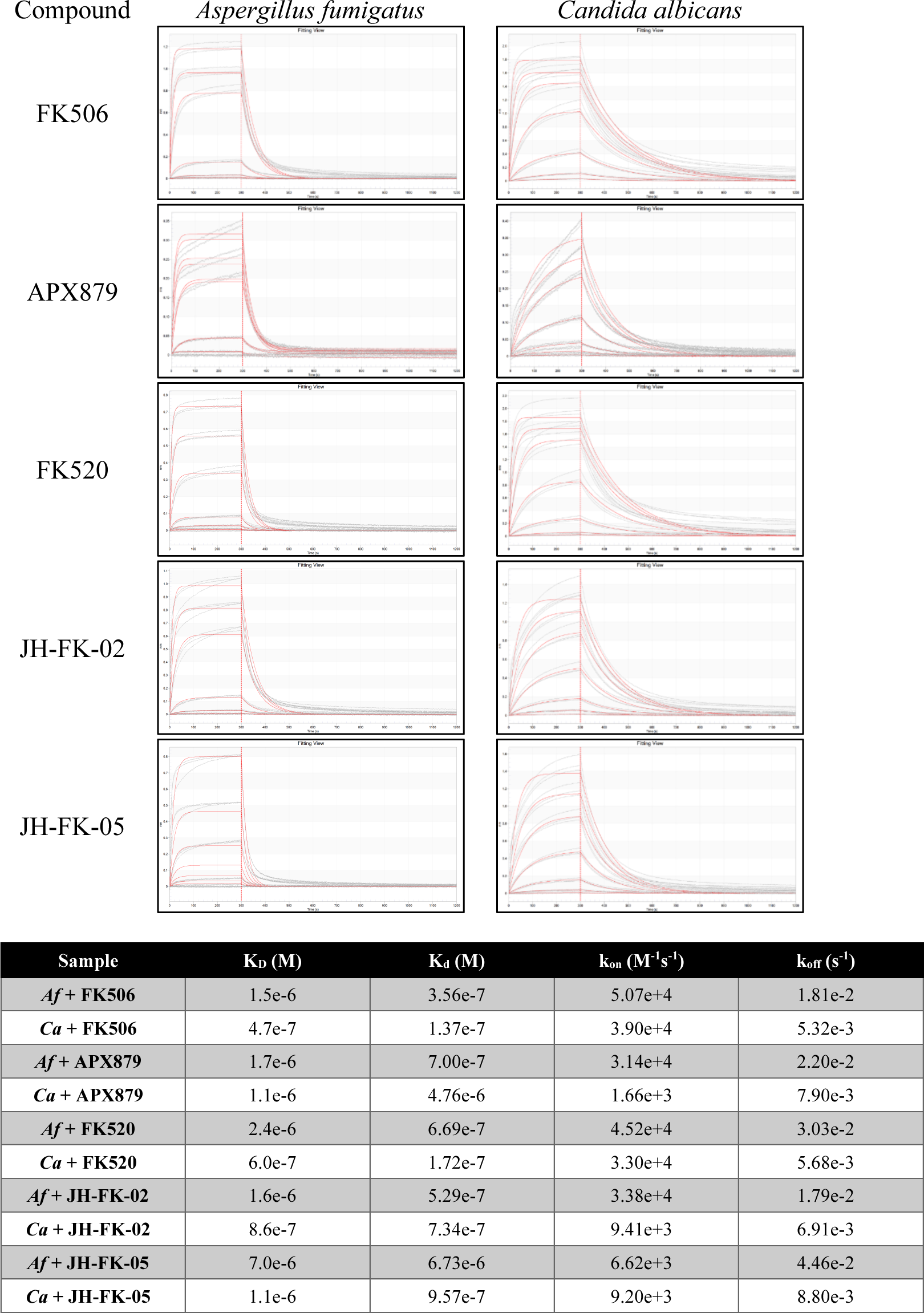
BLI sensorgrams for compound-mediated binding interactions between *A. fumigatus* and *C. albicans* CnA–CnB–calmodulin ternary complex and biotinylated (Avi-tagged)-FKBP12. Experiments were performed using a constant compound concentration of 10 µM and variable analyte protein (CnA–CnB–calmodulin, 2500 – 39.0625 nM by serial 1:1 dilution, plus a 0 nM sample blank). All data are double referenced using sample blanks and parallel reference sensors. Three sample replicates were acquired in each experiment (light gray). Analysis was performed using 1:1 models and global analysis (calculated model traces in red). BLI kinetics and steady state analysis results summarized in table (bottom). Kinetics data were analyzed using 1:1 models and global analysis. Steady state data were analyzed using binding responses in the 290-295 s range of sample association.

**Supplementary Figure 3.**
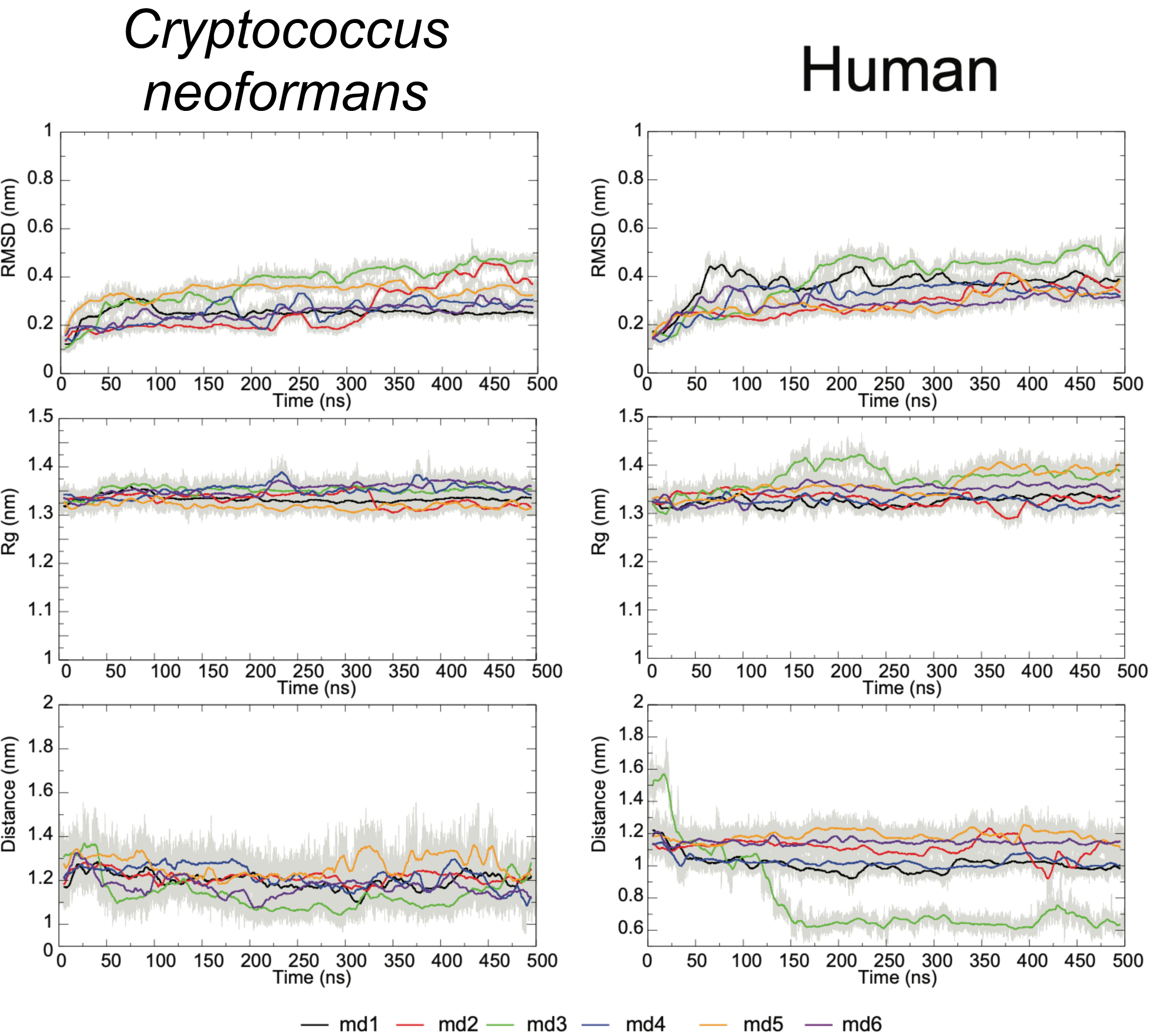
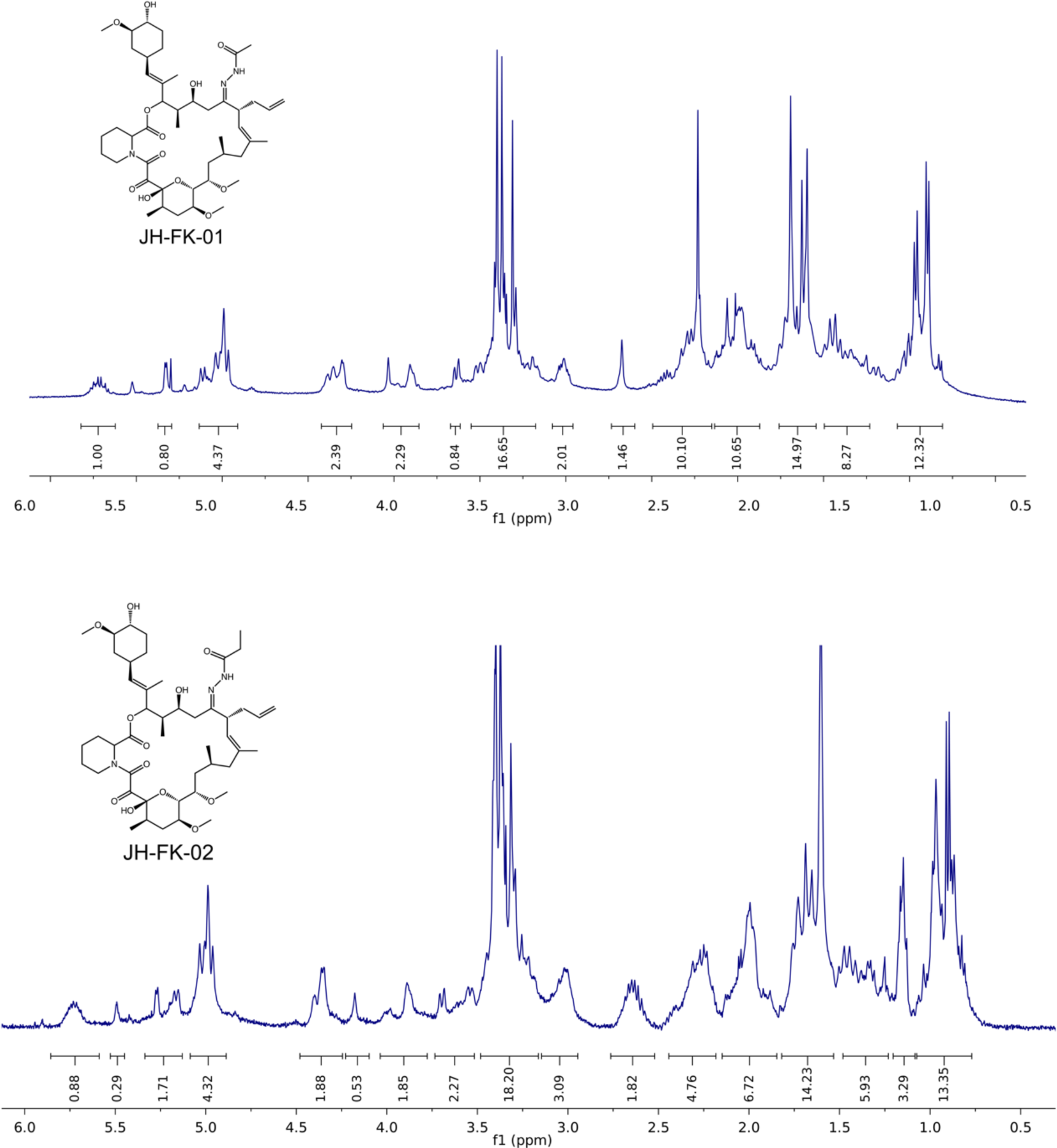
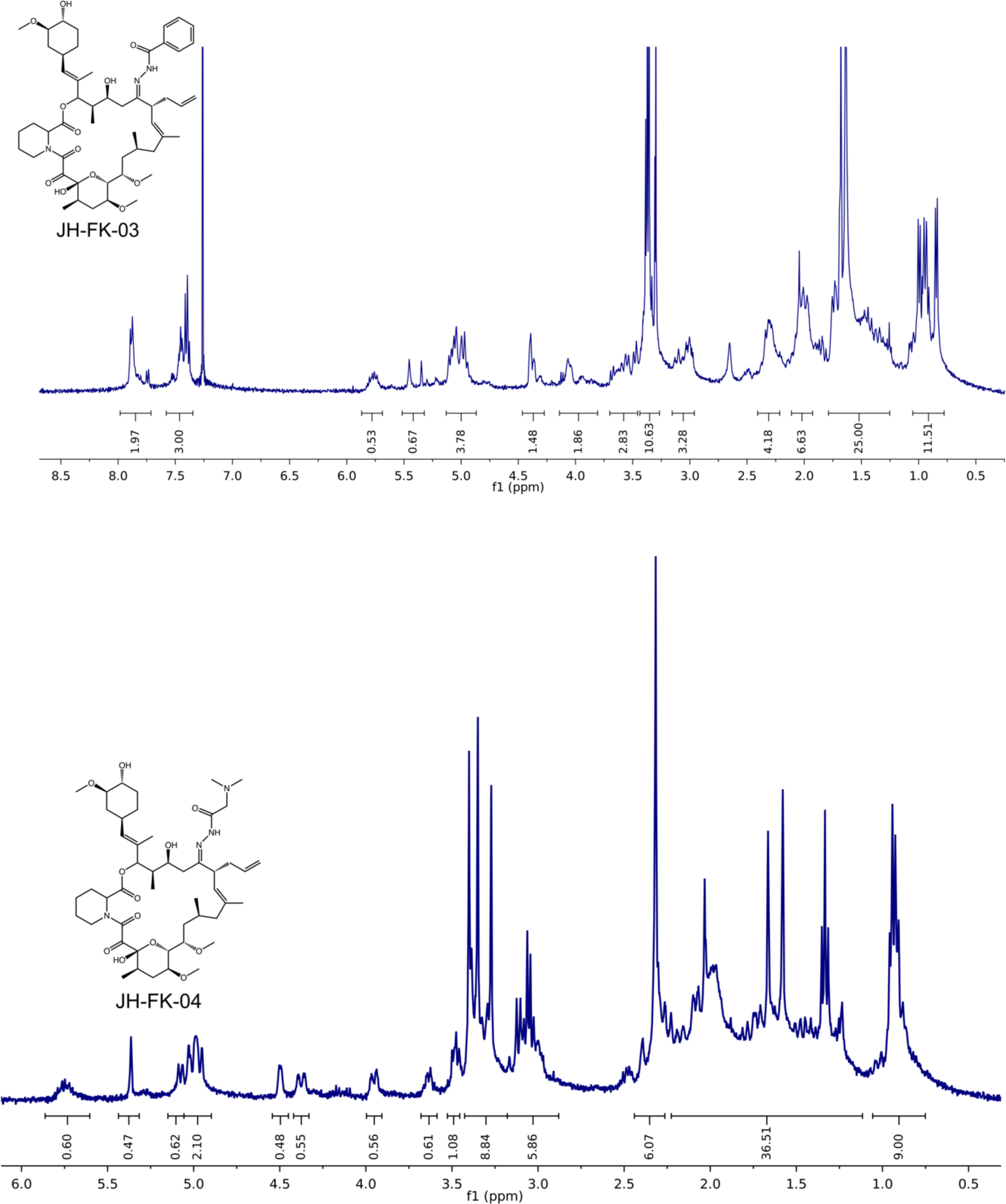
Analysis of 500-ns MD simulation stability. MD simulations were run for JH-FK-05 binding either human FKBP12 or *C. neoformans* FKBP12. Top figure plots the Cα-RMSD for each MD simulation over the length of the simulation. Middle figure plots the radius of gyration (Rg) for each MD simulation over the length of the simulation. Bottom figure plots the center of mass (COM—the 3D point of mass balance for each monomer) between the protein and ligand for each MD simulation over the length of the simulation. MD simulation 3 of *h*FKBP12-JHFK05 was removed from analysis due to unstable COM values observed during the length of the simulation.

**Supplementary Figure 4.**
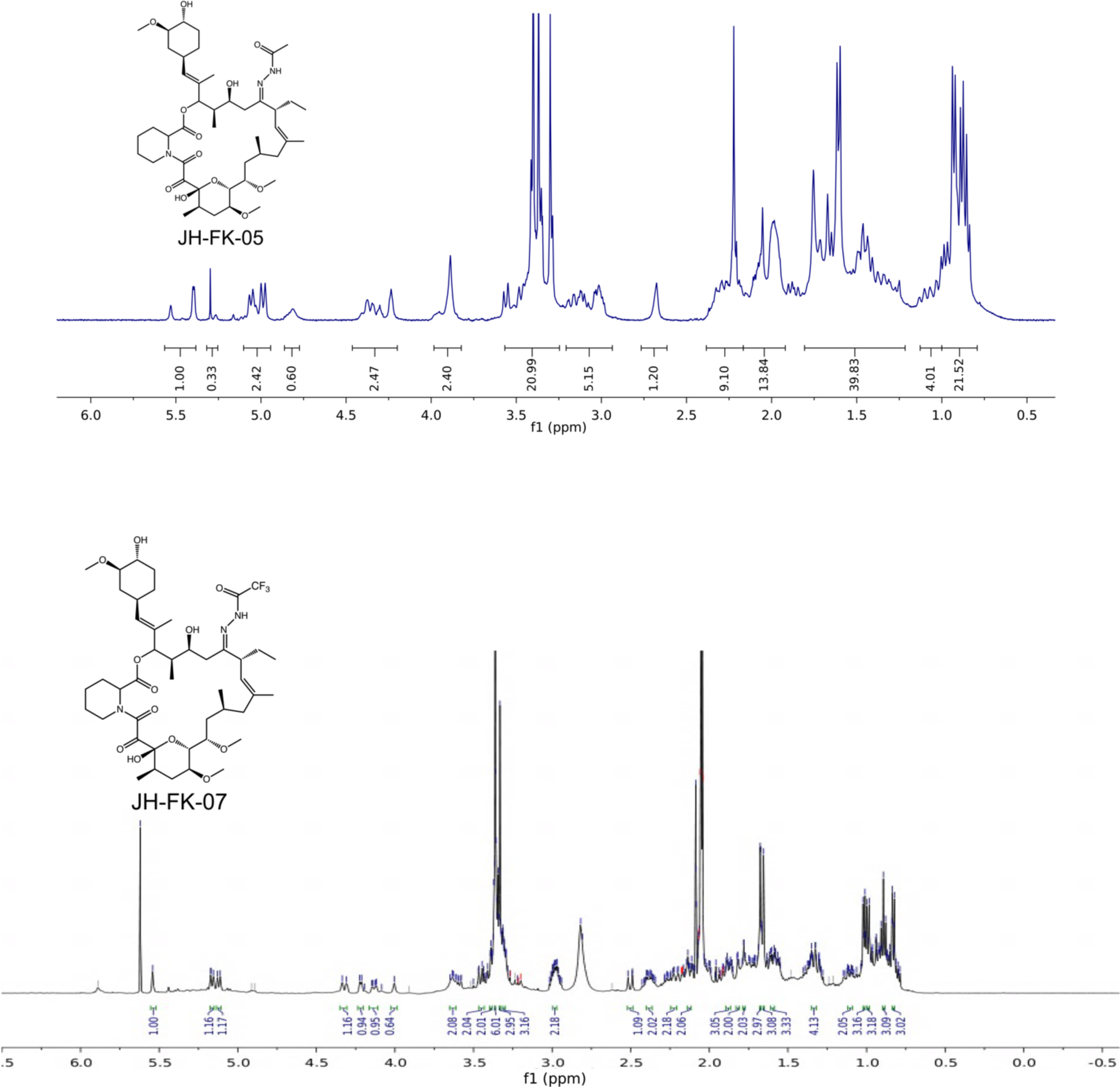
^1^H NMR analysis of FK506/FK520 analogs. All six synthesized FK506/FK520 analogs were subjected to ^1^H NMR analysis. All integrations are indicated. Chemical structure for each compound analyzed is presented at the left.

**Supplementary Table 1.** Crystallographic data and refinement statistics.

| Data name | *Aspergillus fumigatus*<br>FKBP12-FK520<br>PDB 7U0S | *Homo sapiens*<br>miniCN-FKBP12-FK520<br>PDB 7U0T | *Aspergillus fumigatus*<br>miniCN-FKBP12-FK520<br>PDB 7U0U |
| --- | --- | --- | --- |
| **Data collection** |  |  |  |
| Wavelength (Å) | 0.97872 | 0.9786 | 0.9786 |
| Space group | P 2<sub>1</sub> 2<sub>1</sub> 2 | C 1 2 1 | I 2 2 2 |
| **Cell dimensions** |  |  |  |
| a, b, c (Å) | 89.81 91.47 35.53 | 156.1 33.3 64.35 | 75.01 83.7 137.58 |
| α, β, γ (°) | 90.0 90.0 90.0 | 90.0 103.8 90.0 | 90.0 90.0 90.0 |
| Resolution (Å) | 50 - 1.70 (1.74 - 1.70)<sup>a</sup> | 50 - 2.45 (2.51 - 2.45) | 50 - 1.90 (1.95 - 1.90) |
| No. of unique reflections | 32,881 (2,347) | 11,961 (757) | 34,390 (2,485) |
| R<sub>merge</sub> (%) | 6.6 (53.0) | 7.1 (37.0) | 4.8 (57.0) |
| I/σ(I) | 13.78 (2.90) | 13.22 (2.64) | 19.94 (2.34) |
| Completeness (%) | 99.5 (99.2) | 97.8 (85.0) | 99.6 (98.5) |
| Redundancy | 4.8 (4.9) | 3.5 (2.7) | 5.9 (4.6) |
| CC½ | 99.8 (88.2) | 99.7 (83.6) | 99.9 (76.5) |
| **Refinement** |  |  |  |
| Resolution (Å) | 50 - 1.70 | 50 - 2.45 | 50 - 1.90 |
| No. of unique reflections | 32,871 | 11,959 | 34,387 |
| R<sub>work</sub>/R<sub>free</sub><sup>b</sup> | 0.168 (0.233)<br>0.192 (0.269) | 0.203 (0.274)<br>0.259 (0.338) | 0.169 (0.246)<br>0.199 (0.261) |
| **No. of atoms** |  |  |  |
| Protein | 1,845 | 2,274 | 2,415 |
| Ligand | 194 | 64 | 79 |
| Solvent/Water | 227 | 62 | 174 |
| **B factors (Å<sup>2</sup>) (overall)** |  |  |  |
| Protein | 23.56 | 51.35 | 43.17 |
| Ligand | 29.24 | 31.23 | 32.12 |
| Solvent/Water | 39.25 | 38.81 | 41.99 |
| **RMSD** |  |  |  |
| Bond lengths (Å) | 0.006 | 0.002 | 0.006 |
| Bond angles (°) | 0.94 | 0.59 | 0.85 |
| Ramachandran favored (%) | 97.84 | 97.94 | 98.34 |
| Ramachandran allowed (%) | 2.16 | 2.06 | 1.66 |
| Ramachandran outliers (%) | 0 | 0 | 0 |
| Rotamer outliers (%) | 1.02 | 1.26 | 1.52 |
| RMSD, root-mean-square deviation. |  |  |  |
| <sup>a</sup>Highest-resolution shell shown in parentheses. |  |  |  |
| <sup>b</sup>R<sub>free</sub> was calculated with 10% of the data |  |  |  |

**Supplementary Table 2.** High-resolution mass spectrometry (HRMS) data for FK506/FK520 analogs.

|  | Elemental<br>Composition | Calculated<br>Mass | Observed<br>Mass | Error<br>(ppm) | Ion |
| --- | --- | --- | --- | --- | --- |
| JH-FK-01 | $C_{46}H_{73}N_3O_{12}$ | 860.5267 | 860.5276 | -1 | $[M+H]^+$ |
| | | 882.5087 | 882.5091 | -0.5 | $[M+Na]^+$ |
| JH-FK-02 | $C_{47}H_{75}N_3O_{12}$ | 874.5424 | 874.5429 | -0.6 | $[M+H]^+$ |
| | | 896.5243 | 896.5247 | -0.4 | $[M+Na]^+$ |
| JH-FK-03 | $C_{51}H_{75}N_3O_{12}$ | 922.5424 | 922.5416 | 0.8 | $[M+H]^+$ |
| | | 944.5243 | 944.5236 | 0.8 | $[M+Na]^+$ |
| JH-FK-04 | $C_{48}H_{78}N_4O_{12}$ | 903.5689 | 903.5697 | -0.9 | $[M+H]^+$ |
| | | 925.5509 | 925.5507 | 0.1 | $[M+Na]^+$ |
| JH-FK-05 | $C_{45}H_{73}N_3O_{12}$ | 848.5267 | 848.5273 | -0.7 | $[M+H]^+$ |
| | | 870.5087 | 870.5089 | -0.3 | $[M+Na]^+$ |
| JH-FK-07 | $C_{45}H_{70}F_3N_3O_{12}$ | 902.0592 | 902.498 | 0.5 | $[M+H]^+$ |
| | | 924.4804 | 924.4792 | 1.3 | $[M+Na]^+$ |

**Supplementary Table 3.** Antifungal activity of JH-FK-05 against molds.

| Species | Strain | MIC/MEC<br>($\mu\text{g/mL}$) |
| --- | --- | --- |
| *Aspergillus fumigatus* (Wild-type) | (akuBKu80) | 1 |
| *Aspergillus fumigatus* | (akuBKu80<br>hFKBP12) | >8 |
| <u>Azole-Resistant Strains</u> |  |  |
| *Aspergillus fumigatus* | F12776 | 2 |
| *Aspergillus fumigatus* | F14946 | 1 |
| *Aspergillus fumigatus* | F16314 | 1 |
| *Aspergillus fumigatus* | F16216 | 1 |
| *Aspergillus fumigatus* | F7075 | 2 |
| <u>Echinocandin-Resistant Strain</u> |  |  |
| *Aspergillus fumigatus* | EMFR-S678P | 2 |
| <u>Other Aspergillus Species</u> |  |  |
| *Aspergillus calidoustus* | WT | >8 |
| *Aspergillus flavus* | WT | 1 |
| *Aspergillus niger* | WT | <1 |
| *Aspergillus terreus* | WT | 1 |
| <u>Other molds</u> |  |  |
| *Fusarium solani* | WT | >8 |
| *Mucor circinelloides* | WT | 1 |
| *Paecilomyces variotti* | WT | >8 |
| *Rhizomucor pusilus* | WT | >8 |
| *Rhizopus oryzae* | WT | 1 |
| *Scedosporium apiospermum* | WT | >8 |
| *Scedosporium prolificans* | WT | >8 |

**Supplementary Table 4.** Atom index for JH-FK-05 MD analysis.

| Index | Type | Atoms | Index | Type | Atoms |
| --- | --- | --- | --- | --- | --- |
| 1 | C | C01 | 31 | CH | C30/H24 |
| 2 | CH | C02/H02 | 32 | CH | C31/H25 |
| 3 | CH<sub>2</sub> | C03/H03/H44 | 33 | CH<sub>2</sub> | C32/H26/H53 |
| 4 | CH<sub>2</sub> | C04/H04/H45 | 34 | CH<sub>2</sub> | C33/H27/H54 |
| 5 | CH<sub>2</sub> | C05/H05/H46 | 35 | CH<sub>3</sub> | C34/H28/H55/H66 |
| 6 | CH<sub>2</sub> | C06/H06/H47 | 36 | CH<sub>3</sub> | C35/H29/H56/H67 |
| 7 | N | N45 | 37 | CH<sub>3</sub> | C36/H30/H57/H68 |
| 8 | C | C07 | 38 | CH<sub>2</sub> | C37/H31/H58 |
| 9 | C | C08 | 39 | CH<sub>3</sub> | C38/H32/H59/H33 |
| 10 | C | C09 | 40 | CH<sub>3</sub> | C40/H34/H60/H69 |
| 11 | CH | C10/H07 | 41 | CH<sub>3</sub> | C41/H35/H61/H70 |
| 12 | CH<sub>2</sub> | C11/H08/H48 | 42 | CH<sub>3</sub> | C42/H36/H62/H71 |
| 13 | CH | C12/H09 | 43 | CH<sub>3</sub> | C43/H37/H63/H72 |
| 14 | CH | C13/H10 | 44 | CH<sub>3</sub> | C44/H38/H64/H73 |
| 15 | CH | C14/H11 | 45 | C | C59 |
| 16 | CH<sub>2</sub> | C15/H12/H49 | 46 | CH<sub>3</sub> | C60/H01/H43/H65 |
| 17 | CH | C16/H13 | 47 | N | N54 |
| 18 | CH<sub>2</sub> | C17/H14/H50 | 48 | NH | N55/H40 |
| 19 | C | C18 | 49 | O | O46 |
| 20 | CH | C19/H15 | 50 | O | O47 |
| 21 | CH | C20/H16 | 51 | O | O48 |
| 22 | C | C21 | 52 | O | O49 |
| 23 | CH<sub>2</sub> | C22/H17/H51 | 53 | O | O50 |
| 24 | CH | C23/H18 | 54 | OH | O51/H39 |
| 25 | CH | C24/H19 | 55 | O | O52 |
| 26 | CH | C25/H20 | 56 | O | O53 |
| 27 | C | C26 | 57 | OH | O56/H41 |
| 28 | CH | C27/H21 | 58 | O | O57 |
| 29 | CH | C28/H22 | 59 | OH | O58/H42 |
| 30 | CH<sub>2</sub> | C29/H23/H52 | 60 | O | O61 |

