## Supplemental Methods for "Structure-guided synthesis of FK506 and FK520 analogs with increased selectivity exhibit in vivo therapeutic efficacy against Cryptococcus"


**List of Authors**

Michael J. Hoy,[1] Eunchong Park*,[2] Hyunji Lee*,[3†] Won Young Lim*,[3]
D. Christopher Cole,[4] Nicholas D. DeBouver,[5,6] Benjamin G. Bobay,[7]
Phillip G. Pierce,[5,6] David Fox III,[5,6] Maria Ciofani,[2] Praveen R. Juvvadi,[4‡] William Steinbach,[4‡] Jiyong Hong,[3] and Joseph Heitman[1]#

**Affiliations**

1 Department of Molecular Genetics and Microbiology, Duke University Medical Center, Durham, North Carolina, USA

2 Department of Immunology, Duke University Medical Center, Durham, North Carolina, USA

3 Department of Chemistry, Duke University, Durham, North Carolina, USA

4 Division of Pediatric Infectious Diseases, Department of Pediatrics, Duke University Medical Center, Durham, North Carolina, USA

5 UCB Pharma, 7869 NE Day Road West, Bainbridge Island, Washington, USA

6 Seattle Structural Genomics Center for Infectious Disease (SSGCID), Seattle, Washington, USA

7 Duke University NMR Center, Duke University Medical Center, Durham, North Carolina, USA

*=equal contributions

†Current address: Department of Chemistry and the Howard Hughes Medical Institute, University of Illinois at Urbana-Champaign, Urbana, Illinois, USA

‡ Current address: Department of Pediatrics, University of Arkansas for Medical Sciences, Little Rock, Arkansas, USA


Running Head: Novel FK506 analogs increased for fungal specificity

### Methods of Protein Production

#### ***Aspergillus fumigatus* and *Candida albicans* calcineurin A, calcineurin B, and calmodulin complexes**

*C. albicans* calcineurin subunit A with an N-term MGSSHHHHHHSSGENLYFQGS tag (Uniprot Q5ABP4 residues 54 – 609, CaalA.00174.a.EX11, CID11764), *C. albicans* calcineurin subunit B with no tag (Uniprot C4YS24 residues 1 – 173, CaalA.01011.a.EX11), and *C. albicans* calmodulin with no tag (Uniprot P23286 residues 1 – 149, CaalA.00464.a.EX11, CID11763) were separately codon-optimized by ATUM for insect expression, flanked with unique SapI sites (with bases TGATAA prior to the 3' SapI site) and cloned by ATUM via SapI into a customized version of pBacgus-1 (Millipore Sigma, formerly Novagen) which includes two SapI sites within the MCS. One *E. coli* TOP10 transformant was fully sequence-verified over the Open Reading Frame (ORF) and flanking regions and DNA from this clone subjected to SF9 transfection via Bestbac protocols.

*A. fumigatus* calcineurin subunit A with an N-term MGSSHHHHHHSSGENLYFQGS tag (Uniprot Q4WUR1 residues 2 – 434, CID11448, AsfuA.00174.a.FW11), *A. fumigatus* subunit B with no tag (Uniprot Q4WDF2 residues 21 – 193, CID11452, AsfuA.01011.a.FX11), and *A. fumigatus/ Coccidioides immitis* calmodulin (100% identity) with no tag (Uniprot J3KLB7 residues 1 – 149, CID11453, CoimA.00464.a.FX11) were separately codon-optimized by ATUM for insect expression, flanked with unique BamHI/HindIII sites (with TGATAA prior to the HindIII) and intermediate plasmids delivered in a kanamycin-resistant shuttle vector. Genes were released via BamHI/HindIII digestion and ligated into BamHI/HindIII-digested and gel-purified insect transfer vector pFastbac-1 (Life Technologies). A resulting TOP10 transformant for each construct was miniprepped and DNA sequenced across the ORF and flanking regions. Upon validation this DNA was subsequently transposed into DH10Bac cells (Life Technologies) and screened by PCR to identify clones with the desired gene insert. Bacmid DNA was prepped via standard protocols (Life Technologies) from one clone positive for insert by PCR for each construct and the bacmid DNA subjected to SF9 transfection via Bac-to-Bac protocols (Life Technologies).

Calcineurin A, calcineurin B and calmodulin complexes were expressed and purified using similar methods to those previously described (34). In brief, for each species of complex, calcineurin A, calcineurin B and calmodulin were co-transfected into Tni insect cells grown in ESF921 medium (Expression Systems) at roughly a 1:1:1 mixture of viruses such that the final multiplicity of infection (MOI) was 2-3. Cultures were harvested 48-72 h post-infection. Cells were centrifuged and stored at -80 °C. Cell pastes were lysed via hypotonic lysis in 25 mM Tris pH 8.0, 0.02% CHAPS, and protease inhibitor tablets, EDTA-free (Roche), and clarified before applying them to two 5 mL $Ni^{2+}$ charged HiTrap Chelating HP (GE Healthcare) columns. The complex of His-tagged calcineurin A, calcineurin B and calmodulin was eluted with a gradient of 25 mM Tris, pH 8.0, 200 mM NaCl, 1 mM TCEP, and 500 mM Imidazole. Fractions containing the complex were pooled and diluted in buffer to lower the NaCl concentration down to 50 mM, then injected onto HiTrap SP-FF and HiTrap Q-FF (GE Healthcare) ion exchange columns arranged in tandem. After loading and washing in 25 mM Tris pH 8.0, 50 mM NaCl, 1 mM TCEP and 5 mM $CaCl_2$, the columns were separated and eluted using a gradient up to 1 M NaCl. The cleanest protein complex eluted from the HiTrap Q-FF column and was pooled and concentrated to 10-15 mg/mL. Each complex was then injected onto a 120 mL Sephacryl S-300 column (GE Healthcare) equilibrated in 25 mM HEPES, pH 8.0, 50 mM NaCl, 0.5 mM TCEP, and 5 mM $CaCl_2$. Fractions containing the complex in 1:1:1 stoichiometry were pooled and concentrated to 10 mg/mL, then flash frozen in liquid nitrogen and stored at -80 °C (CaalA.00174.a.EX11 [calcineurin A], CaalA.01011.a.EX11 [calcineurin B], CaalA.00464.a.EX11 [calmodulin], Batch ID - PD00426; AsfuA.00174.a.FW11 [calcineurin A], AsfuA.01011.a.FX11 [calcineurin B], CoimA.00464.a.FX11 [calmodulin], Batch ID – PD00411).

**Biotinylated *A. fumigatus* FKBP12 P90G and *H. sapiens* FKBP12**

*A. fumigatus* FKBP12 with N-term MSGSHHHHHHHHGGENLYFQGSGLNDIFEAQKIEWHEGSSGSS tag (Uniprot Q4WLV6 residues 2 – 112 with P90G, AsfuA.18272.a.TM11) and *Homo sapiens*

FKBP12 with N-term MSGSHHHHHHHHGGENLYFQGSGLNDIFEAQKIEWHEGSSGSS tag (Uniprot P62942 residues 2 – 108, HosaA.18272.a.TM11) were separately codon-optimized for *E. coli* expression by Genscript, flanked with unique NdeI/XhoI (including stop codons TGATAA prior to XhoI) and delivered in an ampicillin-resistant shuttle vector. The inserts were released with NdeI/XhoI and ligated into NdeI/XhoI-digested and gel-purified pET28a (modified with an NdeI site replacing NcoI). Following transformation of the ligation one *E. coli* TOP10, one clone per construct was fully sequence-verified over the ORF and flanking regions and subsequently transformed into BL21(DE3) for expression studies.

Bio-*Af*FKBP12 and Bio-*Hs*FKBP12 were each co-transformed with BirA in *E. coli* BL21 (DE3) cells and expression was induced with 1 mM IPTG (GoldBio). Cultures were grown for 16 h at 25 °C then harvested by centrifugation (Beckman) at 10,000 x g for 15 min and pellets stored at -80 °C. Cells were lysed by sonication (Misonix) in 25 mM Tris pH 8.0, 200 mM NaCl, 50 mM L-Arginine monohydrochloride, 0.02% CHAPS (3-[(3-Cholamidopropyl)dimethylammonio]-1-propanesulfonate), 1 mM TCEP (tris(2-carboxyethyl)phosphine) (VWR), 100 mg Lysozyme, 250 U Benzonase and 1 protease inhibitor tablet, EDTA free (Ethylenediaminetetraacetic acid) (Roche). Lysate was clarified by centrifugation at 42,000 RPM (Beckman) for 45 min at 4 °C and filtered with a 0.2-0.8 µm gradient filter. The supernatant was applied to a 5 mL $Ni^{2+}$ charged HiTrap chelating HP (GE Healthcare) columns and the protein was eluted by step gradient of 25 mM Tris pH 8.0, 200 mM NaCl, 1 mM TCEP and 300 mM imidazole. Fractions containing FKBP12 were pooled and digested with TEV protease and dialyzed (3.5 kDa MWCO dialysis cassettes) into 25 mM Tris pH 8.0, 200 mM NaCl, 1 mM TCEP overnight at 4 °C to remove the N-terminal 8x histidine (confirmed by SDS-PAGE). The dialyzed and TEV cleaved target was then re-applied to a 5 mL $Ni^{2+}$ charged HiTrap chelating HP and FKBP12 was collected in the flow-through and early wash fractions. Again, fractions containing FKBP12 were pooled and dialyzed into the final buffer containing 25 mM Tris pH 8.0, 200 mM NaCl, 1mM TCEP. The final samples were concentrated to 18.2 mg/mL (Bio-*Af*FKBP12) and 7.1 mg/mL (Bio-*Hs*FKBP12) and flash frozen in liquid nitrogen and stored at -80 °C (AsfuA.18272.a.TM11, Batch ID - PD38408; HosaA.18272.a.TM11, Batch ID – PD38410).

***H. sapiens* and *A. fumigatus* truncated calcineurin A (calcineurin B binding helix) – calcineurin B fusion constructs**

*A. fumigatus* calcineurin subunit A – subunit B fusion with N-term MSGSHHHHHHHHGGENLYFQGS tag (Uniprot Q4WUR1 residues 335 – 370 fused with Uniprot Q4WDF2 residues 1 – 193 on either side of a GGGSSGGSTSGGSSGGG residue linker, AsfuA.00174.a.TQ11 + AsfuA.01011.a.TR11), and *Homo sapiens* calcineurin subunit A – subunit B fusion with MSGSHHHHHHHHGGENLYFQGS tag (Uniprot Q08209 residues 337 – 372 fused with Uniprot P63098 2 -170 on either side of a GGGSSGGSTSGGSSGGG linker, HosaA.00174.a.TU11 + HosaA.01011.a.TV11), were cloned as discussed for FKBP12.

*Af*CnA-CnB or *Hs*CnA-CnB were transformed individually into *E. coli* BL21 (DE3) cells and expression was induced with 1mM IPTG (GoldBio) at 18 °C or 37 °C, respectively. The culture was grown for 16 h and harvested by centrifugation (Beckman) at ~10,000 *g* for 15 min. Pellets were stored at -80 °C. Cells were lysed and purified using methods described for *Af*FKBP12 through the secondary Ni-chromatography step, of which fractions containing each fusion were pooled and concentrated for SEC (*Hs*CnA-CnB at 14.4 mg/mL, *Af*CnA-CnB at 3.3 mg/mL). Each complex was loaded onto a Superdex 75 16/60 PG column and eluted in 25 mM Tris pH 8.0, 200 mM NaCl, 1 mM TCEP, and 5 mM $CaCl_2$. Final sample was concentrated to 10.61 mg/mL (*Hs*CnA-CnB) and 11.3 mg/mL (*Af*CnA-CnB), flash frozen in liquid nitrogen and stored at -80 °C. (AsfuA.00174.a.TQ11+AsfuA.01011.a.TR11 fusion, Batch ID – PD38406; HosaA.00174.a.TU11+HosaA.01011.a.TV11 fusion, Batch ID – PD38411)
